# Tandem association of CLOCK:BMAL1 complexes on DNA enables recruitment of CBP/p300 through multivalent interactions

**DOI:** 10.1101/2025.09.04.674342

**Authors:** Diksha Sharma, Lisa Stoos, Megan R. Torgrimson, Priya Crosby, Kyle M. Franks, Nicole C. Parsley, Miles Membreno, Georg Kempf, Lukas Kater, Chelsea L. Gustafson, Hsiau-Wei Lee, Seth M. Rubin, Alicia K. Michael, Nicolas H. Thomä, Carrie L. Partch

## Abstract

The basic helix-loop-helix PER-ARNT-SIM (bHLH-PAS) transcription factor CLOCK:BMAL1 interacts with E-box motifs in the context of nucleosomes to elicit a circadian pattern of gene expression that oscillates with approximately 24 hour periodicity. Core clock genes and other highly rhythmic targets of CLOCK:BMAL1 typically possess a tandem arrangement of E-boxes that is required for robust oscillations. Here, we show that the presence of tandem E-boxes enables CLOCK:BMAL1 to bind more internal sites on the nucleosome, leading to release of DNA from the histone core and the presentation of multiple coactivator binding motifs in close proximity to facilitate multivalent interactions with the coactivator CBP/p300. We show that the transactivation domain (TAD) of BMAL1, essential for CLOCK:BMAL1 activity, interacts with several modular domains of CBP. Deletion of these CBP domains or chemical inhibition of protein-protein interactions with CBP significantly reduces or eliminates CLOCK:BMAL1-driven activity. Altogether, this suggests that multivalent interactions with CBP may play a role in the ability of tandem CLOCK:BMAL1 heterodimers to recruit this limiting cofactor in cells.

## Introduction

The transcription factor CLOCK:BMAL1 sits at the core of several interlocking transcription-translation feedback loops that generate mammalian circadian rhythms^1^. CLOCK:BMAL1 binds thousands of sites throughout the genome with a preference for E-box motifs; approximately 1400 of these are co-occupied by all of the core clock proteins at some point throughout the day^2^. Highly rhythmic transcriptional targets of CLOCK:BMAL1, including the core clock genes, have tandem E-boxes with a precise spacing of 6 or 7 nucleotides between them that is essential for robust circadian oscillations^3, 4^. We recently showed how both CLOCK and BMAL1 use their PAS domains to interact with each other and the histone octamer to permit occupancy at tandem E-boxes on the outer edge of the nucleosome^5^.

CLOCK:BMAL1 activity relies on two elements within the intrinsically disordered tails that follow the PAS domain core of the heterodimer. First, the 51-residue coiled-coil domain of CLOCK encoded by exon 19 homodimerizes^6^ and contributes to cooperative binding at tandem E-box motifs^7^. Its deletion in the *Clock^Δ19^* mouse gives rise to dominant negative behavior that diminishes activation of E-box-driven transcription and leads to low amplitude or disrupted circadian rhythms *in vivo*^8^. Second, the transactivation domain (TAD) at the C-terminus of BMAL1 is essential for CLOCK:BMAL1 activity^9^ and circadian rhythms *in vivo*^10^. The BMAL1 TAD binds to coactivators and repressors at overlapping sites^11–14^ to establish the paradigm that competition for the BMAL1 TAD leads to oscillating gene expression throughout the day. While the molecular basis for CLOCK:BMAL1 repression by cryptochromes has been fairly well studied^13, 15–18^, there are still questions about how the BMAL1 TAD interacts with CBP/p300^2, 19, 20^ to allow CLOCK:BMAL1 to effectively compete for these pleiotropic coactivators^21^ in cells.

CBP and p300 have several small modular domains and intrinsically disordered motifs flanking the acetyltransferase core that interact directly with transcription factors (TFs), some of which employ multivalent interactions with CBP/p300 to enhance recruitment to their target genes^22–24^. We previously determined that the BMAL1 TAD interacts with the KIX domain of CBP and p300, identifying mutations in the conserved IXXLL coactivator binding motif of the TAD that eliminate CLOCK:BMAL1 activity and disrupt cellular circadian rhythms^13^. However, the most rhythmic transcriptional targets of CLOCK:BMAL1, including the core clock genes, possess tandem E-boxes that are essential for oscillatory gene expression^3, 4^. As binding to tandem E-boxes would allow the presentation of multiple TADs in close spatial proximity, we wanted to test if the BMAL1 TAD interacts with other TF-recruitment domains on CBP/p300.

Here, we employed SeEN-seq^25^ to explore CLOCK:BMAL1 binding to multiple E-box motifs on the nucleosome. Strikingly, we discovered that an unfavorable single internal E-box becomes a viable target for CLOCK:BMAL1 when present as a tandem array spaced by 6 or 7 nucleotides. Using NMR spectroscopy, we show that two regions of the BMAL1 TAD each bind one of the well-studied TF binding sites on the CBP KIX domain: the TAD helix binds the c-Myb site, while the TAD switch region binds to the MLL1 site. Additionally, we found that both TAD regions are also involved in binding to CBP TAZ1 and TAZ2 domains. Deletions of TAZ1 and/or KIX significantly reduce CBP coactivation of CLOCK:BMAL1 on a *Per1-*luciferase reporter, while blocking access to endogenous CBP KIX with small molecules inhibits CLOCK:BMAL1 transactivation. Together, this demonstrates the potential importance of multivalent interactions with CBP/p300 for CLOCK:BMAL1 function, which is facilitated by its binding to tandem E-box motifs.

## Results

### CLOCK:BMAL1 binds to internal tandem E-box motifs on the nucleosome

The Widom 601 DNA sequence contains a fixed E-box motif (CACGTG) at superhelical location (SHL) +5.1. However, the structured core of the CLOCK:BMAL1 heterodimer, comprising its basic helix-loop-helix and two PAS domains (bHLH-PAS-AB), is unable to bind this internal site^5^. To systematically investigate the spatial constraints and preferred spacing for CLOCK:BMAL1 binding to double E-box motifs within a nucleosome, we tiled a second E-box across the Widom 601 sequence, creating a double E-box library, and performed SeEN-seq on reconstituted nucleosomes (Fig. 1a). Comparison with a prior SeEN-seq profile from a single E-box library^5^, generated using the same tiled E-box but on a Widom 601 background lacking the fixed E-box at SHL +5.1, revealed nearly identical enrichment on the left side of the nucleosome (SHL –7 to –5), consistent with the two E-boxes being too far apart for cooperative binding (Fig. 1b). By contrast, the right side of the profile exhibited new enrichment peaks near SHL +4, where the tiled E-box approaches the fixed one, suggesting a preferred spacing of 3 or 6 nucleotide base pairs (bp) between the two motifs. Additional peaks emerged near SHL +6, where the enrichment had previously dropped in the single E-box context, corresponding to tiled E-boxes located 3 or 7 bp downstream of the fixed motif position. While CLOCK:BMAL1 is known to preferentially bind tandem E-box motifs spaced 6–7 bp apart^3, 4^, a pattern we confirm on nucleosomes, along with an additional 3 bp spacing, we find that the double E-box configuration notably expands CLOCK:BMAL1’s ability to engage more internal positions within the nucleosome.

**Figure 1:**
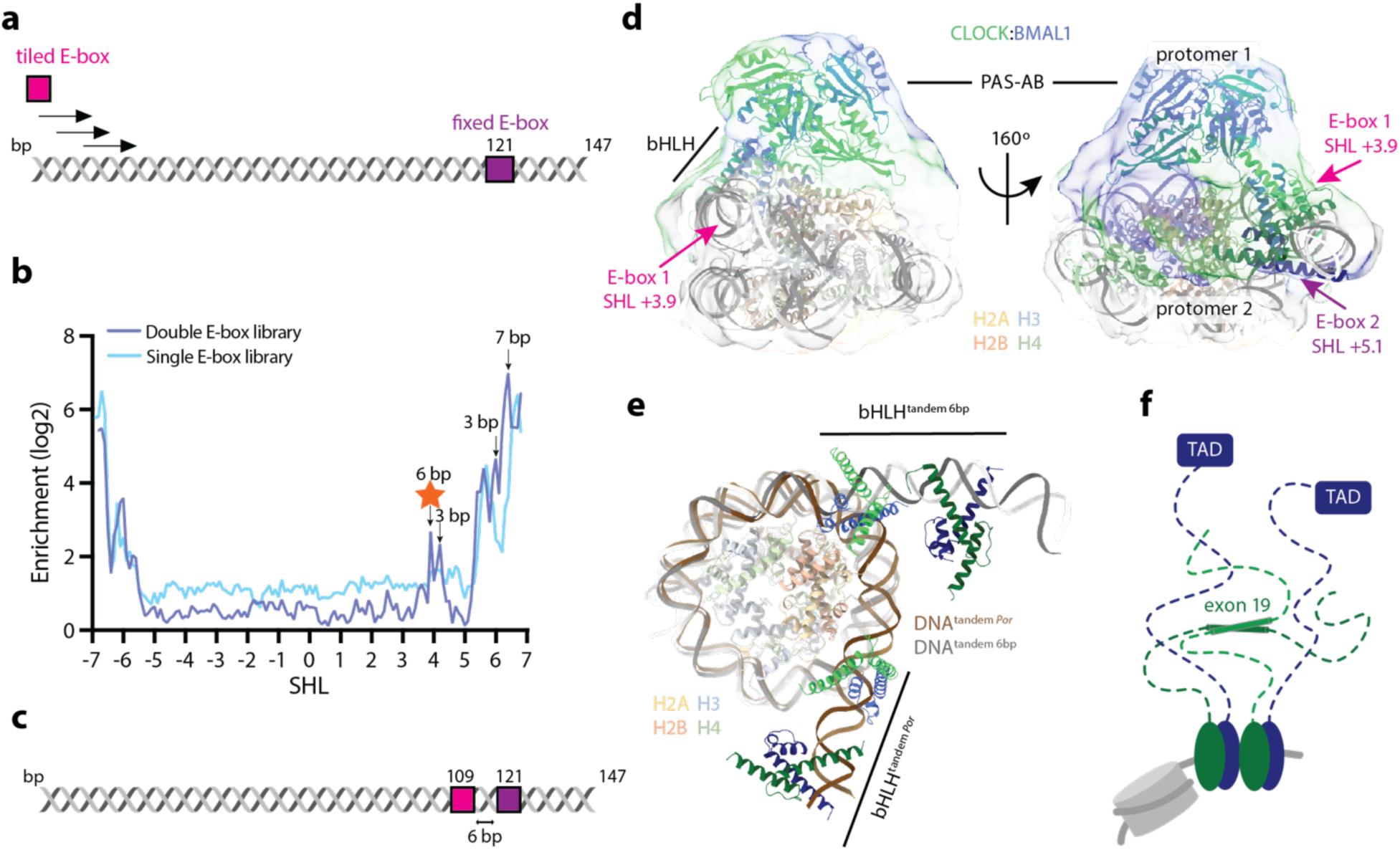
Cooperativity between two CLOCK:BMAL1 protomers enables binding to more internal positions within the NCP. a) Schematic of the double E-box SeEN-seq library used in this study. One E-box was fixed at SHL+5.1 (tile 121) and one E-box was tiled through the W601 sequence to determine the preferred spacing between two CLOCK:BMAL1 protomers *in vitro*. b) Comparison of SeEN-seq profiles of CLOCK:BMAL1 on a single or double E-box library. Values shown are an average from 3 independent replicates. The indicated SHL of the E-box corresponds to the centroid of the motif CACGTG. Arrows indicate preferred spacing between the two E-boxes and the DNA sequence used for structural studies is highlighted (star). c) Schematic of the W601 DNA with tandem E-boxes separated by 6 bp. d) Overlay of the cryo-EM map of two CLOCK:BMAL1 protomers bound to E-box motifs at SHL+3.9 and +5.1 (i.e., end of first E-box at SHL+4.2 and beginning of second E-box at +4.8) with a model containing the two bHLH domains and the PAS domains of the first protomer. e) Comparison of the E-box positions in CLOCK:BMAL1-NCP*^Por^*and CLOCK:BMAL1-NCP^tandem^ ^6bp^. In CLOCK:BMAL1-NCP^tandem^ ^6bp^ the tandem E-boxes are positioned more than 10 bp more internally within the nucleosome. f) Cartoon depicting the disordered C-termini of CLOCK (green) and BMAL1 (blue) with domains that are known to contribute to regulation of CLOCK:BMAL1.

Mass photometry experiments of CLOCK:BMAL1 and a nucleosome containing tandem CACGTG E-boxes with 6 bp spacing at SHL +3.9/+5.1 (tandem 6bp) (Fig. 1c, corresponding to a SeEN-seq peak in Fig. 1b) demonstrated formation of a complex with the expected mass for two heterodimers on the nucleosome, as well as an additional peak suggesting that three CLOCK:BMAL1 protomers can associate with nucleosomes containing internal tandem E-boxes, possibly reflecting flexible PAS-domain interactions (Fig. S1a). To study how CLOCK:BMAL1 accesses these internal sites, a cryo-EM structure of this E-box nucleosome core particle (NCP) template was solved (CLOCK:BMAL1-NCP^tandem^ ^6bp^) (Fig. 1d, Fig. S1 and S2, Table 1) at an overall resolution of 4.1 Å. The structure revealed that two CLOCK:BMAL1 protomers collectively release approximately 10 bp of nucleosomal DNA compared to binding at more external E-box motifs in the native *Por* promoter (Fig. 1e). The protomers interact directly via their PAS domains, although mapping these interactions was limited by low resolution (8-12 Å) in this region. The orientation of the bHLH domain of the inner protomer was determined using cross-linking mass spectrometry (Fig. S1b), while placement of its PAS domains was guided by the electron density. The inner protomer binds with its bHLH domain nearly perpendicular to the nucleosome plane and the CLOCK PAS-B domain positioned atop the acidic patch (Fig. 1d). For the external protomer, only the bHLH domains were confidently modelled as both the electron density and the crosslinks did not allow unambiguous modelling of the PAS domains (Fig. S1c,d). The orientation of the bHLH domains, however, places the PAS domains in proximity to histones H2A and H2B.

Given CLOCK:BMAL1’s preference for tandem E-boxes with both 6 or 7 bp spacing, we attempted complex formation with a nucleosomal template containing the outer E-box fixed at SHL+5.1 and the inner E-box shifted 1 bp further inward to directly compare the effects of 6 vs 7 bp spacing. Upon confirming complex formation by mass photometry (Fig. S1e), we proceeded with solving another cryo-EM structure of this complex (CLOCK:BMAL1-NCP^tandem^ ^7bp^) at 4.2 Å overall resolution (Fig. S1f, Fig. S3, Table 1). While it was possible to distinguish the densities corresponding to each protomer, model building was restricted to the bHLH domains (Fig. S1f). In comparison to CLOCK:BMAL1-NCP^tandem^ ^6bp^, the angle between the two inner bHLH domains differs by 20° and the trajectory of the released DNA by 15° (Fig. S1g,h respectively). Most noticeable, however, is the rearrangement of the PAS domains (compare Fig. S1i and S1j). In CLOCK:BMAL1-NCP^tandem^ ^7bp^, the PAS domains of the inner protomer no longer contact the histone core but instead form an extensive interface with the PAS domains of the outer protomer. These are positioned atop histones H2A, H2B, and H4, while the bHLH domains are located next to the nucleosome.

In addition to the release of DNA by binding to internal motifs and the PAS-PAS contacts, we also noted that binding of two CLOCK:BMAL1 heterodimers in close proximity on DNA could facilitate the activity of two regions in their intrinsically disordered C-termini that are required for activity of the transcription factor, exon 19 of CLOCK and TAD of BMAL1 (Fig. 1f). In particular, we were intrigued by the idea that the presentation of two BMAL1 TADs in close proximity could facilitate interactions with multiple modular TF recruitment domains on CBP/p300.

### Two motifs in the BMAL1 TAD bind the CBP KIX domain

We previously showed that the BMAL1 TAD interacts directly with the KIX domain from CBP and p300^13^. However, the KIX domain harbors two distinct TF binding interfaces that are named for the well-studied TFs that bind there, the c-Myb site and the MLL1 site (Fig. 2a)^26^. In some cases, two TADs within a single TF bind to both sites on the KIX domain, as with p53 and FOXO3a (Fig. S4a)^27, 28^. A previous study suggested that both sites on the KIX domain are involved in binding to the BMAL1 TAD^29^; however, given the presentation of two TADs by tandemly bound CLOCK:BMAL1 on chromatin, we wanted to explore this interaction in more detail.

**Figure 2:**
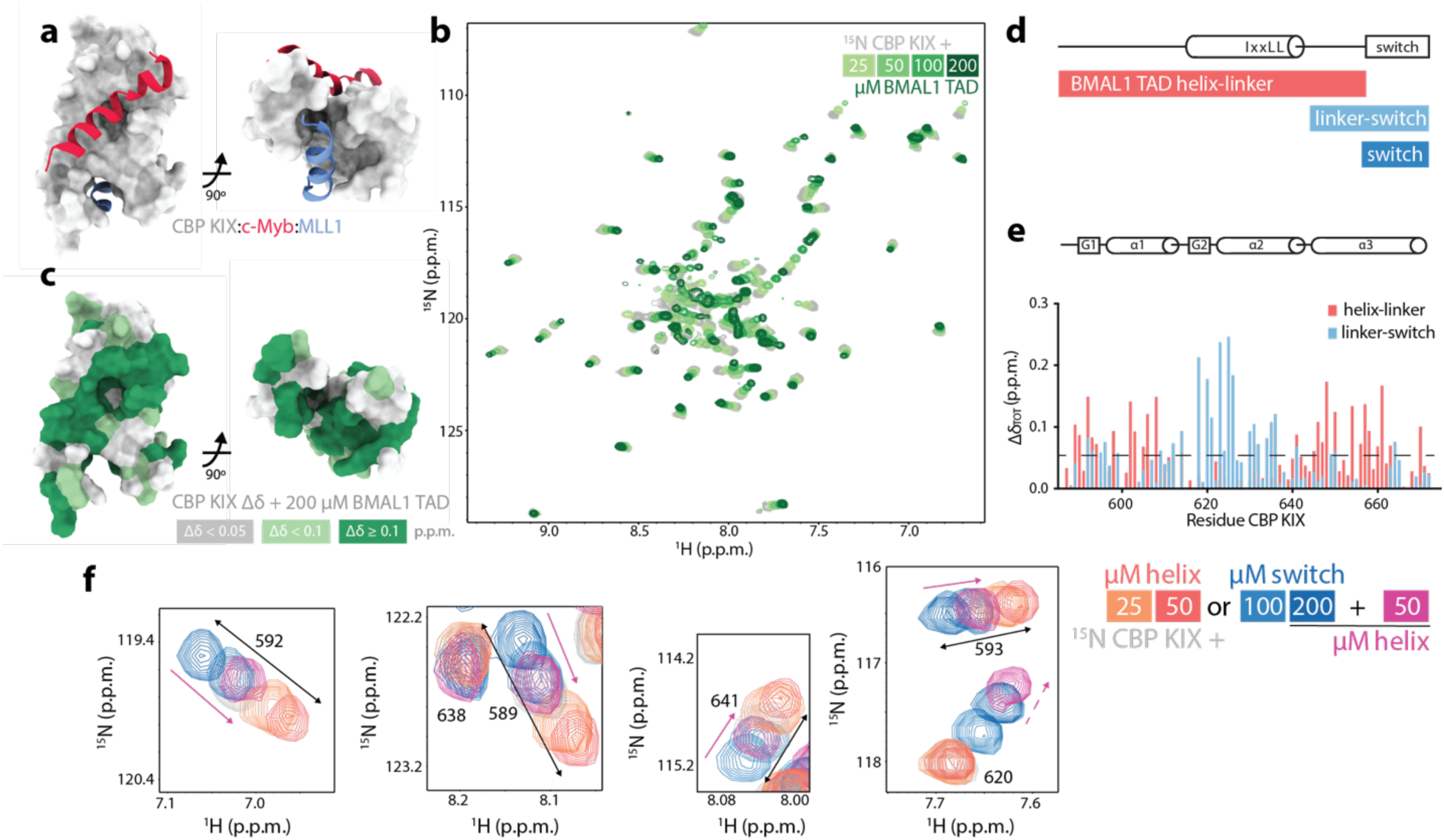
The BMAL1 TAD engages both TF binding sites on the CBP KIX domain. a) CBP KIX domain (gray, PDB 2AGH) with c-Myb (red) and MLL1 (blue) peptides bound. b) ^15^N-^1^H HSQC spectra of 100 µM ^15^N CBP KIX (gray) titrated with BMAL1 TAD as indicated (light to dark green). c) Chemical shift perturbations (CSPs) on ^15^N CBP KIX with 200 µM BMAL1 TAD mapped onto the KIX structure. d) Schematic of the BMAL1 TAD (above) and truncated peptides used (below). e) Secondary structure of CBP KIX (above) and CSPs on ^15^N CBP KIX from the BMAL1 TAD peptides at 100 µM (below). Dashed line, significance cutoff of 0.05 p.p.m. CSPs are mapped onto the KIX structure in Fig. S4d. f) Selected regions of the ^15^N-^1^H HSQC of ^15^N CBP KIX depicting the two-state equilibrium with the BMAL1 TAD helix-linker (salmon), switch (blue), or 200 µM switch and 50 µM helix-linker peptides together (magenta). Solid arrows, addition of helix-linker peptide moves peaks from the switch to helix reporting state; dashed arrow, addition of the helix-linker peptide enhances binding of the switch at the MLL1 binding site.

To determine TAD binding sites on the CBP KIX domain, we employed NMR spectroscopy, monitoring chemical shift perturbations (CSPs) of ^15^N-labeled CBP KIX in the presence of increasing amounts of a natural abundance (i.e., NMR-invisible) BMAL1 TAD construct containing residues 579-626 that encompasses the entire KIX binding site we determined earlier^13^. Figure 2b shows extensive concentration-dependent perturbation of chemical shifts in ^15^N CBP KIX from titration of the BMAL1 TAD, which largely map in and around the two previously described TF binding sites (Fig. 2c and Table 2). Although most of the chemical shift trajectories were linear and displayed fast exchange behavior, we identified a group of residues with chemical shift trajectories that curved at higher concentrations (Fig. S4b), indicating that they are sensitive to multiple binding events. These residues were localized within the core of the KIX domain that is involved in allosteric regulation between the two TF binding sites (Fig. S4c)^26, 30, 31^.

We previously identified two motifs in the BMAL1 TAD that bind to the coactivator CBP/p300 KIX or the repressor CRY1, separated by a glycine/proline-rich linker^13^. These are depicted in Figure 2d as a coactivator-binding IXXLL motif within the partially formed helix and the C-terminal 8 residues known as the switch region, named for the *cis/trans* isomerization of the W624-P625 imide bond^32^. The same two regions were recently identified in BMAL1 by a high-throughput scan of TF motifs capable of both activation and repression^33^, independently confirming their capacity to bind both coactivators and repressors.

To see if these two BMAL1 TAD motifs might target different sites on the KIX domain, we added peptides of either the helix-linker (salmon) or the linker-switch (light blue) into ^15^N CBP KIX and monitored CSPs at 1:1 stoichiometry (Fig. 2e). These data provide evidence that the BMAL1 TAD helix and switch regions bind to largely distinct sites on the KIX domain: the TAD helix induces perturbations primarily at the c-Myb site on KIX helices alpha-1 and alpha-3, and the switch region perturbs the MLL1 site comprised by alpha-2, alpha-3, and the G2 helix (Fig. 2e, Fig. S4d, and Table 2).

Perturbations at sites on the KIX domain by both the helix and switch regions, such as on alpha-1, report on perturbation of the allosteric core of the KIX domain^31^. Here, we observed chemical shift patterns on CBP KIX indicative of a two-state exchange that reports on the apparent occupancy of the c-Myb and MLL1 sites. For example, the TAD helix-linker peptide induced shifts of these KIX domain peaks linearly in one direction, while the switch peptide induced shifts in the opposite direction along the same linear trajectory (Fig. 2f). To test if these shifts respond to changes in TF binding between the two motifs, we first added 200 µM switch peptide to ^15^N CBP KIX (in blue) and then added 50 µM helix peptide (in magenta). We observed that binding of the TAD helical motif to the c-Myb site on KIX shifted the apparent equilibrium of the allosteric core (e.g., residues 589, 592, 641) back towards the helix-bound state, and stimulated binding of the switch peptide at the MLL1 site using residue 620 as a reporter (Fig. 2f). Together, these data suggest that a single BMAL1 TAD occupies both TF binding sites on the CBP KIX domain, consistent with the 1:1 stoichiometry of the complex observed before by isothermal titration calorimetry (ITC)^13^.

### Both *cis* and *trans* isomers of the BMAL1 TAD switch region bind the MLL1 site on KIX

To explore binding of the switch region in more detail, we collected a series of ^15^N-^1^H HSQC spectra titrating the linker-switch peptide or a fragment of the isolated switch comprising residues 619-626 (FSDLPWPL) into ^15^N CBP KIX. Both peptides induced a similar pattern of chemical shift perturbations in the KIX domain (Fig. 3a), with the longer peptide having a modestly higher apparent affinity. This could be due to truncation of the isolated switch peptide at residue F619, which contains a motif used by p53 (i.e., FSDL in its activation domain 1) to bind the MLL1 site on the KIX domain^27^. High concentrations of the isolated switch peptide were required to begin reaching saturation, consistent with a K_D_ for the CBP KIX domain and the isolated TAD switch in the hundreds of micromolar (Fig. 3b).

**Figure 3:**
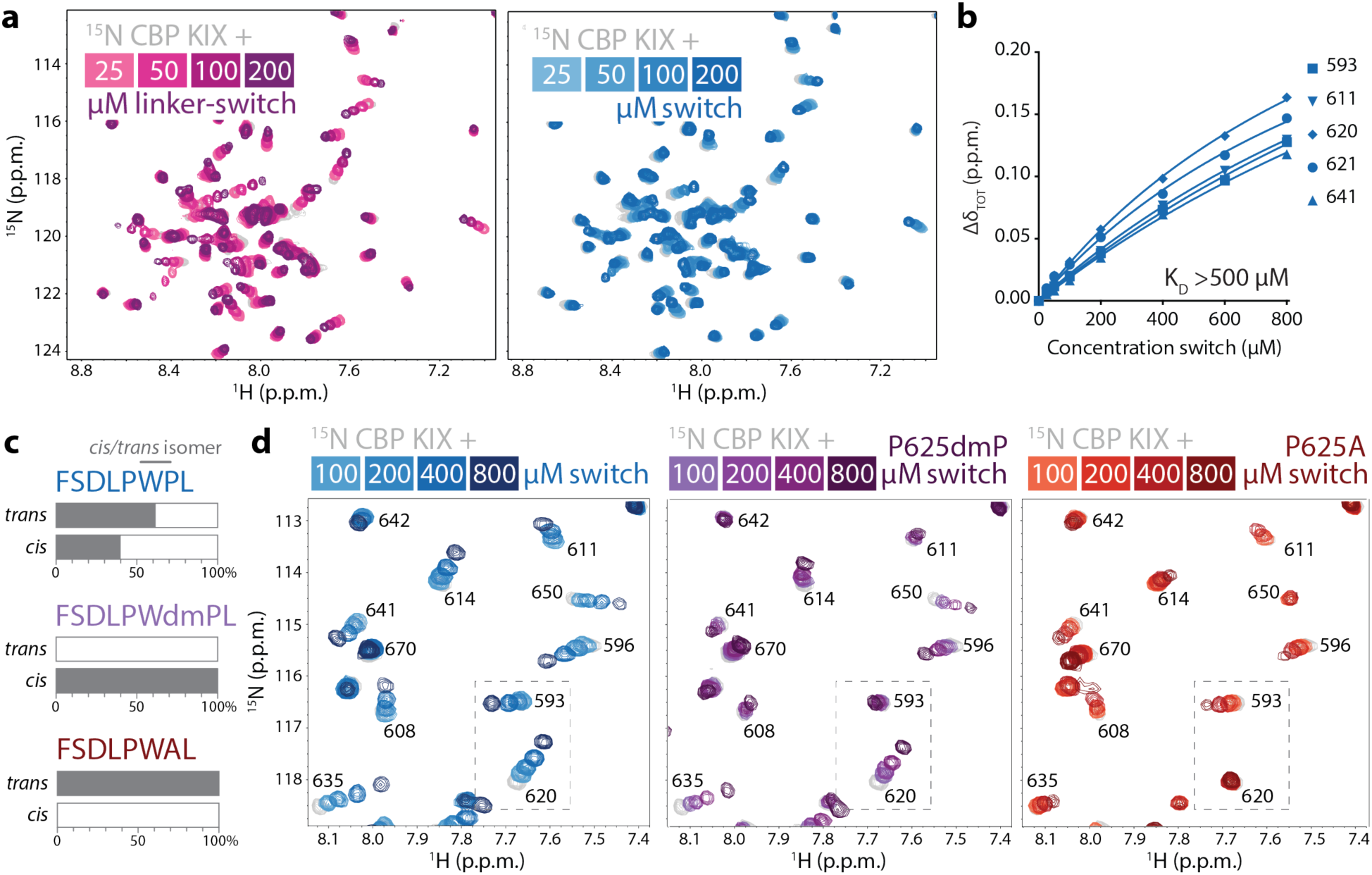
*Cis* and *trans* isomers of the BMAL1 TAD switch both bind to CBP KIX. a) ^15^N-^1^H HSQC of 100 µM ^15^N CBP KIX titrated with BMAL1 linker-switch (light to dark pink) or switch (light to dark blue) peptides. b) Fitting concentration-dependent CSPs on ^15^N CBP KIX numbered residues to estimate a K_D_ for the BMAL1 switch. c) Approximate populations of *cis* and *trans* isomers in switch with mutants; dmP, dimethyl proline. d) ^15^N- ^1^H HSQC of 100 µM ^15^N CBP KIX with switch (blue), *cis*-locked P625dmP-switch (purple), and *trans*-locked P625A switch (red). Dashed box, CSPs that highlight *cis or trans-*specific shifts in ^15^N CBP KIX, both of which are evident in the switch peptide containing both isomers.

The switch region of the BMAL1 TAD has an apparent population of approximately 60% *trans* and 40% *cis* Pro 625 isomer as assessed by NMR^32^. Using dimethyl proline at this position (P625dmP), we created a switch peptide that is effectively locked in an all *cis* conformation, while the P625A substitution generated a switch peptide in all *trans* (Fig. 3c)^32^. Using the native switch peptide and these locked versions, we examined how each bind to the ^15^N CBP KIX domain using NMR spectroscopy. The pattern of chemical shift perturbations collectively demonstrate that both *cis-*locked and *trans*-locked isomers can bind at the MLL1 site (Fig. 3d). For example, residue 614 in the core of the MLL1 site on KIX shifted with titration of the native switch peptide, and to a lesser degree with each of the locked peptides, suggesting that they contribute additively to this shift. However, there was evidence for *cis* or *trans*-specific shifts as depicted in the dashed box in Figure 3d. Some of these perturbations were localized at the MLL1 site (i.e., residue 620) and some affected the allosteric core (residue 593 or 635), demonstrating that the two TAD switch isomers impart distinct effects on the KIX domain. However, longer TAD constructs containing both the TAD helix and these locked versions of the switch bound with approximately the same affinity^32^, suggesting that the helix region dictates the overall affinity of BMAL1 TAD for the CBP KIX domain.

### The primary KIX-TAD binding site occurs between the TAD helix and the c-Myb site

To investigate the importance of BMAL1 TAD binding at the c-Myb site on the KIX domain, we used a well-studied triple mutant comprising the substitutions Y650A, A654Q, and Y658A on CBP KIX (Fig. 4a). This triple mutant (3mut) acts as a loss-of-function mutant for TFs like CREB and c-Myb that rely on this binding site to recruit CBP/p300 for coactivation^34–36^. We examined binding of wild-type (WT) and 3mut CBP KIX to the BMAL1 TAD by ITC, demonstrating that interaction of the mutant KIX is disrupted under conditions in which WT KIX binds with a K_D_ of approximately 2 µM (Fig. 4b)^13^.

**Figure 4:**
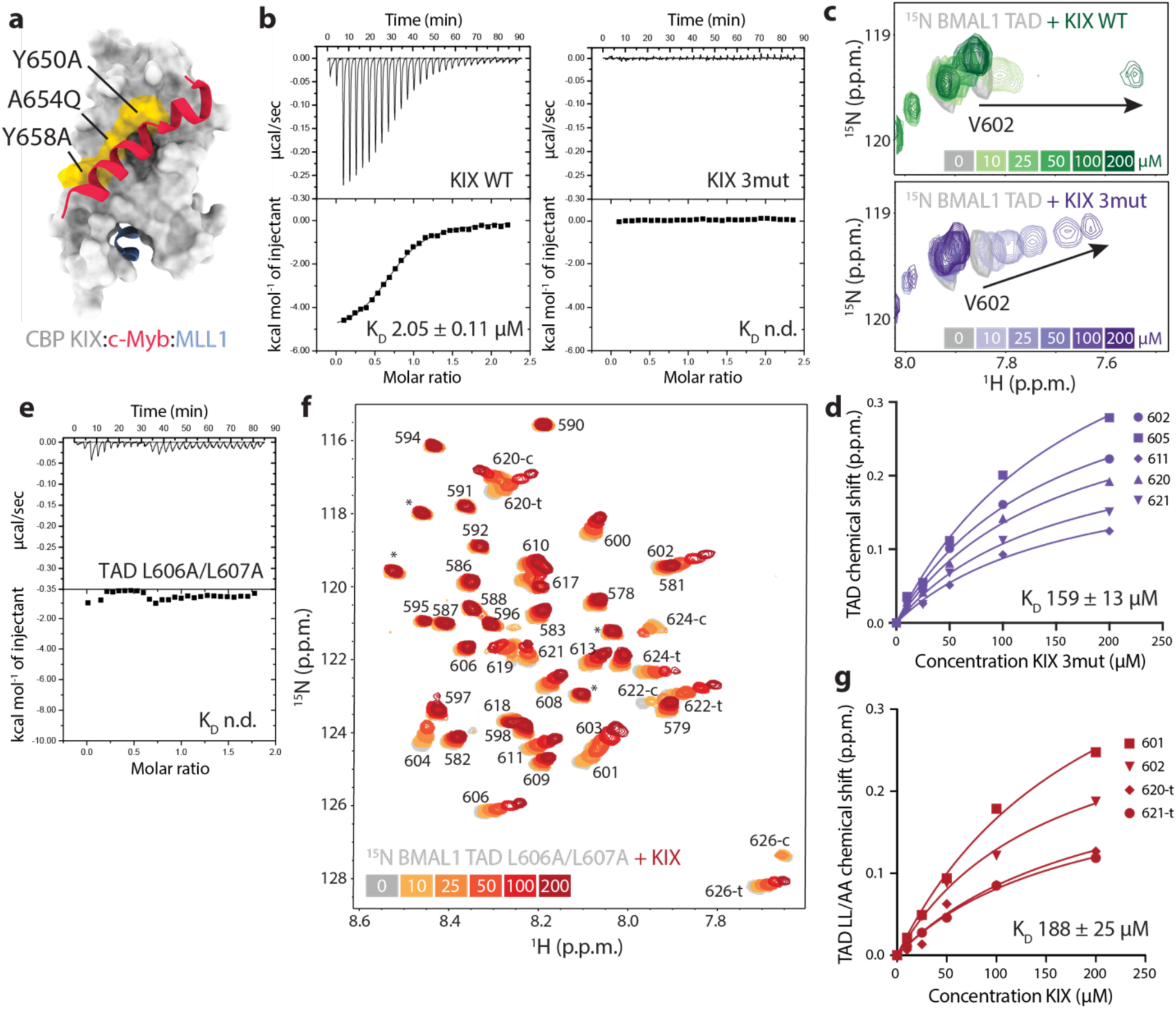
The c-Myb site on CBP KIX is the primary BMAL1 TAD binding site. a) CBP KIX domain (gray, PDB 2AGH) with c-Myb (red) and MLL1 (blue) peptides and the 3mut residues (yellow). b) ITC data of WT CBP KIX (left) or 3mut CBP KIX (right) with the BMAL1 TAD. c) ^15^N-^1^H HSQC of 100 µM ^15^N BMAL1 TAD residue V602 (gray) titrated with WT CBP KIX (top, light to dark green) or 3mut CBP KIX (bottom, light to dark purple). d) Fitting concentration-dependent CSPs on ^15^N BMAL1 TAD to estimate a K_D_ for 3mut CBP KIX. e) ITC data of CBP KIX with the L606A/L607A BMAL1 TAD. f) ^15^N-^1^H HSQC of 100 µM ^15^N BMAL1 TAD L606A/L607A (gray) titrated with CBP KIX (orange to red). Switch peaks (619-626) are labeled c (*cis*) or t (*trans*). g) Fitting concentration-dependent CSPs on ^15^N BMAL1 TAD L606A/L607A numbered residues to estimate a K_D_ for CBP KIX.

NMR is ideal for looking at low affinity interactions, where titration experiments typically display fast exchange behavior exemplified by concentration-dependent peak ‘walking’ between the apo and fully bound states^37^. To examine how the triple mutant affects KIX binding to the TAD, we collected ^15^N-^1^H HSQC spectra of ^15^N BMAL1 TAD in the presence of increasing concentrations of CBP KIX WT (top) or 3mut (bottom, Fig. 4c). Titration with WT KIX gave rise to chemical shift perturbations in the ^15^N TAD in the intermediate exchange regime, consistent with its low micromolar affinity, while titration with the KIX 3mut was in fast exchange, consistent with significantly lower affinity. Fitting the concentration-dependent shifts in ^15^N TAD induced by KIX 3mut gave an approximate K_D_ of 160 µM for the BMAL1 TAD (Fig. 4d).

The BMAL1 TAD helix has an IXXLL motif in its partially formed helix that is a conserved coactivator binding site^38, 39^. We previously showed that disruption of this motif with the L606A/L607A mutation eliminates activation of a *Per1-*luciferase reporter by CLOCK:BMAL1 and fails to generate circadian rhythms when expressed in *Bmal1-/-* PER2::LUC fibroblasts^13^. Measuring binding of the L606A/L607A TAD mutant to WT KIX by ITC under the same conditions as above did not yield any binding heats (Fig. 4e), so we turned to NMR spectroscopy for quantitative information on binding. Upon titrating WT KIX into ^15^N L606A/L607A BMAL1 TAD, we observed concentration-dependent chemical shift perturbations in fast exchange (Fig. 4f). These shifts occurred in residues of both the TAD helix (e.g., residues 601, 602) and switch region (residues 620, 621) and were collectively fit to an apparent K_D_ of approximately 190 µM (Fig. 4g). Therefore, mutations in either the BMAL1 TAD helix or the CBP KIX c-Myb site both reduce the affinity by about 100-fold, suggesting that this is the primary interface for recruitment of the KIX domain by the BMAL1 TAD.

### Multiple TF-binding domains of CBP interact directly with the BMAL1 TAD

Given that one BMAL1 TAD fully occupies the CBP/p300 KIX domain, we wondered if the TAD could also interact with other CBP/p300 domains, which would enhance its recruitment to CLOCK:BMAL1 at tandem E-boxes. CBP and p300 are large proteins with at least six TF-binding modular domains or disordered motifs that flank the catalytic core represented by the central bromodomain through the histone acetyltransferase (HAT) domain (Fig. 5a). We used a [5,6] TAMRA-labeled BMAL1 peptide comprising a minimal TAD (residues 594-626) with helix and switch regions that are ∼99% identical in sequence from humans to honeybees^13^ to measure binding to isolated TAZ1, KIX, and TAZ2 domains using fluorescence polarization (FP) (Fig. 5b). This revealed that the TAD can bind to all 3 domains with the rank order in affinity of TAZ1>KIX>TAZ2 (high to low affinity). Many TFs bind to more than one of these domains in CBP/p300^40^, and it is particularly common for TFs that bind DNA as oligomers and present multiple TADs, like the p53 tetramer^23^.

**Figure 5:**
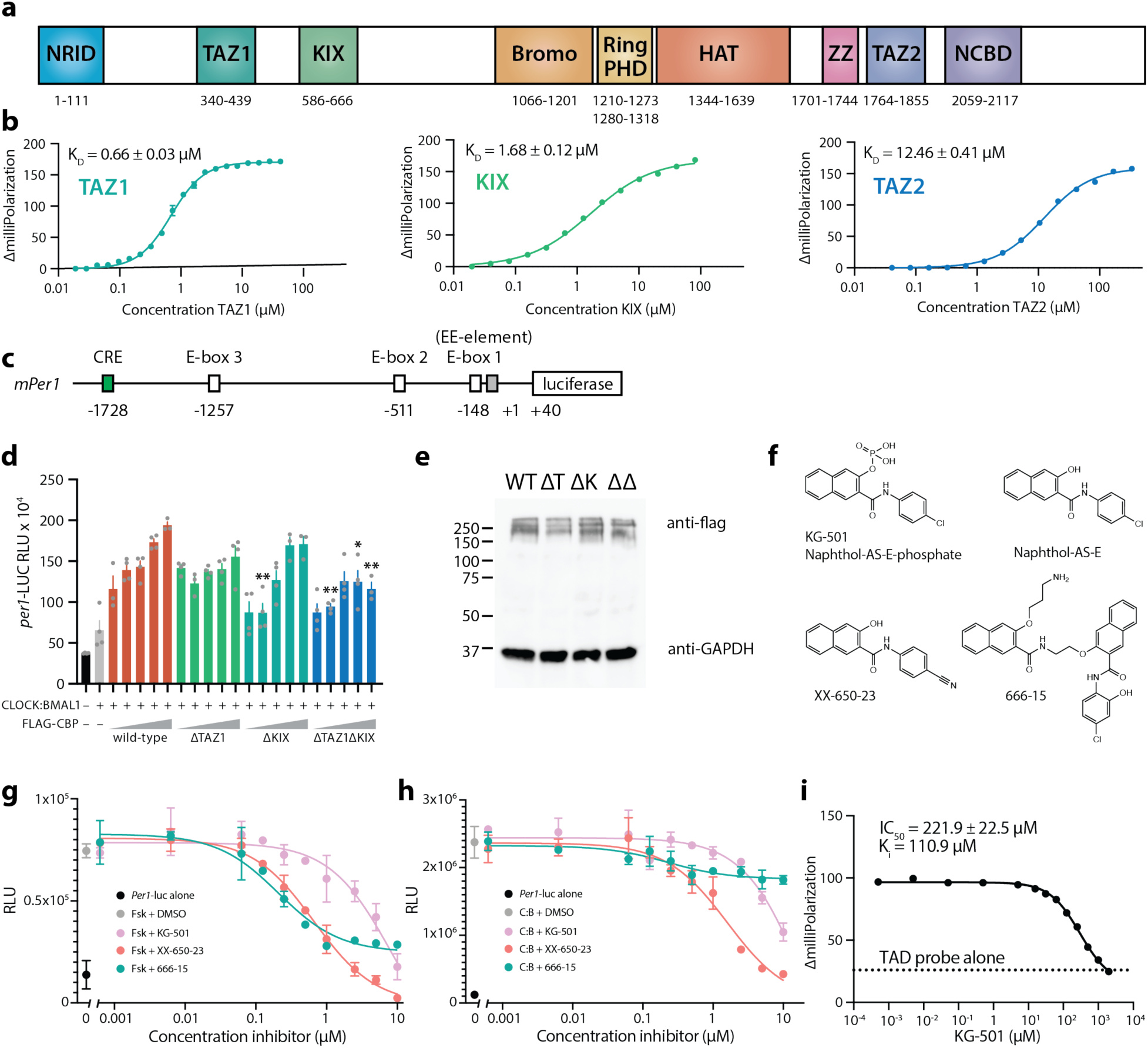
Multiple TF recruitment domains on CBP bind to the BMAL1 TAD for multivalent regulation of CLOCK:BMAL. a) Schematic of CBP domains (mouse CBP numbering, UniProt P45481). b) Fluorescence polarization data of CBP TAZ1 (left), CBP KIX (middle), or CBP TAZ2 (right) binding to 25 nM [5,6] TAMRA-BMAL1 TAD probe. n = 3, mean ± s.d. c) Schematic of mouse *Per1-*luciferase with CRE and E-boxes, including the E-box1 and a degenerate E-box (gray) that comprise the EE-element. Numbering listed below are base pair positions relative to the transcriptional start site. d) *Per1*-luciferase reporter assay in HEK293T cells transfected with CLOCK:BMAL1 (gray) and increasing concentrations of Flag-tagged WT CBP (orange), CBP ΔTAZ1 (green), CBP ΔKIX (teal), or CBP ΔTAZ1ΔKIX (blue). n = 4, mean ± s.d., representative of 4 independent experiments. Tukey’s multiple comparisons test vs WT at the same concentration. Only significant differences are indicated: *, p≤0.05; **, p≤0.01. e) Western blot of CBP WT and deletion mutants expressed in HEK293T cells with GAPDH used as loading control. n = 2. f) Structures of CREB inhibitors. g) Activation of *Per1*-luc reporter in HEK293T cells after forskolin (Fsk) stimulation in the presence of DMSO or inhibitors. n = 3, mean ± s.d. h) Activation of *Per1*-luc reporter in HEK293T cells after transfection with CLOCK:BMAL1 (C:B) in the presence of DMSO or inhibitors. n = 3, mean ± s.d. i) Fluorescence polarization with 2 µM CBP KIX bound to 10 nM [5,6] TAMRA-labeled minimal BMAL1 TAD probe and increasing concentrations of KG-501. IC_50_ reported is the mean from 2 replicates done in duplicate and K_i_ calculated from the Cheng-Prusoff equation.

To test how multivalent interactions could affect coactivation of CLOCK:BMAL1 by CBP, we looked at activation of a mouse *Per1*-luciferase reporter in human embryonic kidney HEK293T cells. This 2 kb fragment of the promoter from the mouse *Per1 gene* contains a CREB-response element (CRE) and three E-boxes, the most proximal of which to the transcription start site occurs as a tandem E-box with a degenerate E-box separated by 6 bp (Fig. 5c)^3, 8^. The area around this so-called tandem EE-element is responsible for most of the activity from this reporter^8, 41^. We transfected HEK293T cells with the *Per1-*luc reporter, *Clock, Bmal1,* and either empty vector or increasing amounts of CBP wild-type (WT) or the domain deletions as indicated (Fig. 5d). Under these conditions, CBP WT elicited an approximate 4-fold increase in CLOCK:BMAL1-driven reporter activity. Deletion of the TAZ1 or KIX domains alone subtly blunted the dose-dependent response, although this was not consistently statistically significant in this assay. Only deletion of both TΑZ1 and KIX together significantly and consistently reduced CLOCK:BMAL1 activation, even though the CBP variants were all expressed to comparable levels (Fig. 5e). These data demonstrate that multiple domains of CBP contribute to CLOCK:BMAL1 activation.

### Small molecule inhibition of the CBP KIX domain blocks CLOCK:BMAL1 activation

To explore the potential role of multivalency from a different perspective, we looked at CLOCK:BMAL1 activation of the *Per1*-luc reporter in HEK293T cells in the presence of compounds that selectively target the endogenous CBP KIX domain. KG-501 (naphthol-AS-E-phosphate) was identified as a c-Myb site binder in an NMR screen of the CBP KIX domain that led to dose-dependent inhibition of forskolin-induced gene expression by CREB (Fig. 5f)^42^. The dephosphorylated version^43^ and a related substituent (XX-650-23)^44^ are more active in cells, although they suffer from poor solubility. Molecule 666-15 is another CREB inhibitor that works independently of the CBP KIX domain^45^. First, we confirmed inhibition of CREB-induced *Per1*-luc activation^46^ by treating HEK293T cells transiently expressing the reporter with the adenyl cyclase activator forskolin, which activates CREB, in the presence of DMSO or increasing amounts of the inhibitors (Fig. 5g). Each of the inhibitors attenuated reporter activation, with KG-501 having moderately decreased potency relative to the others as reported earlier^43^. We then looked at CLOCK:BMAL1 activity after transfecting HEK293T cells with plasmids for *Clock*, *Bmal1*, and the reporter, treating them with DMSO or the set of inhibitors. In contrast to CREB regulation, only the molecules that target the c-Myb site of the KIX domain inhibited CLOCK:BMAL1 activation (Fig. 5h). Finally, we demonstrated direct competition of KG-501 with BMAL1 TAD for the KIX domain. Increasing concentrations of KG-501 displaced the CBP KIX domain in an FP assay with the [5,6] TAMRA-labeled minimal TAD, with a K_i_ similar to that reported for the displacement of phospho-CREB from KIX^42^ (Fig. 5i). These data show that chemically blocking the c-Myb site on the KIX domain of endogenous CBP is sufficient to disrupt multivalent recruitment of CBP/p300 and impinge on coactivation of CLOCK:BMAL1 at E-boxes.

### BMAL1 TAD binding to CBP TAZ1 and TAZ2 utilizes both helix and switch regions

To see if the BMAL1 TAD uses the same helix and switch motifs to bind the CBP TAZ domains, we acquired ^15^N -^1^H HSQC spectra of ^15^N BMAL1 TAD in the presence of increasing concentrations of CBP TAZ1 (Fig. 6a) or TAZ2 (Fig. 6c). We observed a significant broadening of TAD peaks upon addition of TAZ1, although the loss of peak intensity was not uniform across the protein. Figure 6b depicts the differential broadening across the TAD sequence with addition of TAZ1, in which the helix and switch regions exhibited the most severe broadening. TAZ2 binds the BMAL1 TAD with approximately 20-fold lower affinity than TAZ1 (Fig. 5). When we titrated CBP TAZ2 into ^15^N BMAL1 TAD and monitored chemical shift perturbations, we observed two major changes (Fig. 6d). First, the peaks for residues 601-609, comprising the MAV**I**MS**LL**EA sequence (IXXLL) within the TAD helix, immediately broadened. At equimolar concentrations of TAZ2 and TAD, several new peaks appeared, most likely representing this helical motif in its bound state. Second, the remaining chemical shift perturbations mapped throughout the rest of the TAD shifted with apparent fast exchange behavior (Fig. 6d). Unlike other regulators of the BMAL1 TAD studied to date by NMR^11–14^, both TAZ domains influence chemical shifts in an isoleucine and aspartate-rich region upstream of the TAD helix (Fig. 6b, d) with similarity to a SUMO-interaction motif^47, 48^. The [5,6] TAMRA-labeled minimal TAD peptide we used to assess binding by FP lacks this N-terminal motif, so the apparent K_D_s we determined for the TAZ domains (Fig. 5b) likely underrepresent their actual affinity.

**Figure 6:**
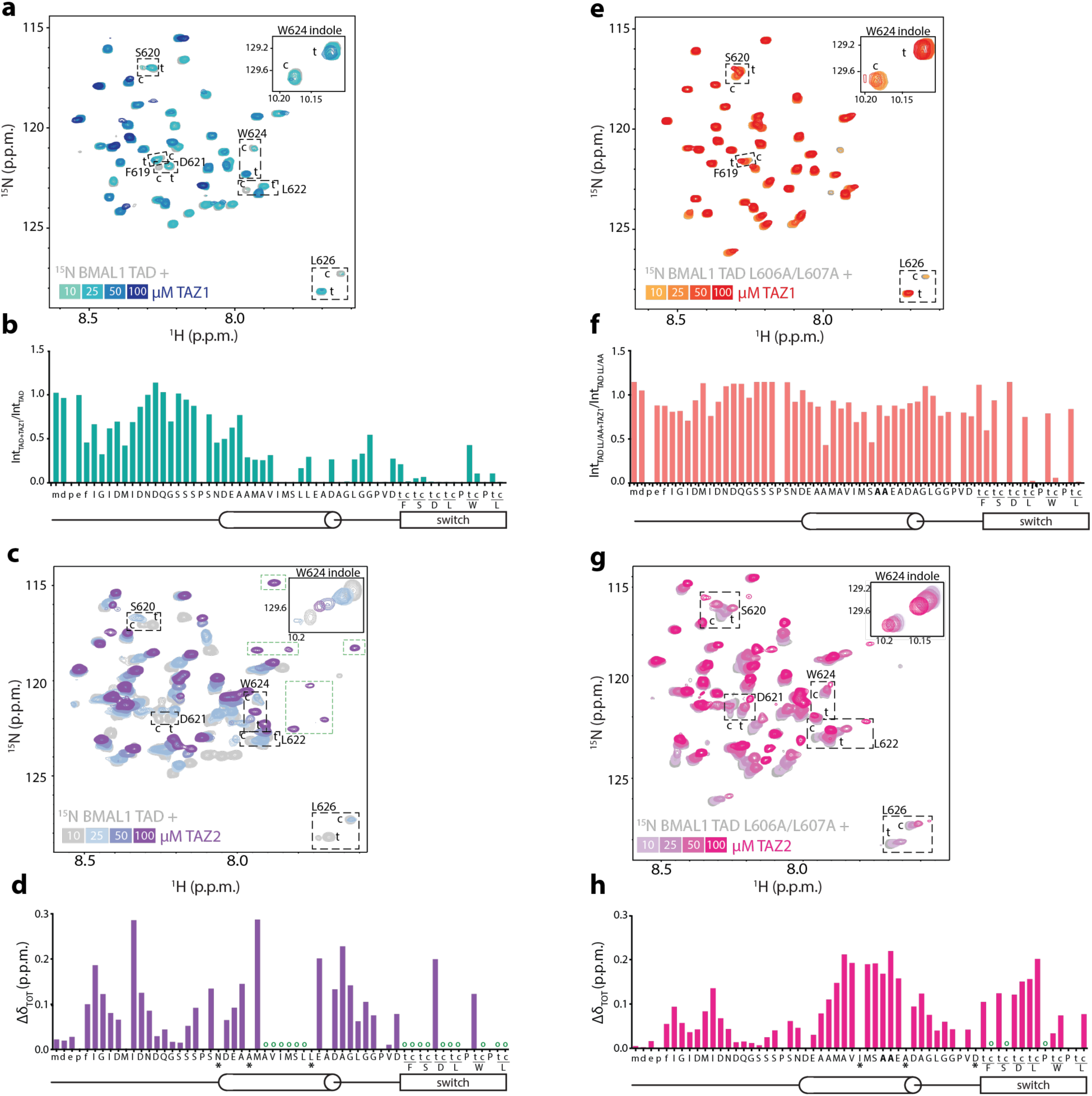
CBP TAZ domains bind to the helix and switch regions of the BMAL1 TAD. a) ^15^N-^1^H HSQC of 100 µM ^15^N BMAL1 TAD (gray) titrated with CBP TAZ1 (teal to dark blue). Dashed boxes, switch residues with c (*cis*) and t (*trans*) peaks. Inset, W624 indole peak. b) Differential broadening of 100 µM ^15^N BMAL1 TAD in the presence of 50 µM CBP TAZ1. Both peaks from the switch (c, t) are plotted. c) ^15^N-^1^H HSQC of 100 µM ^15^N BMAL1 TAD (gray) titrated with CBP TAZ2 (light to dark purple). d) CSPs of 100 µM ^15^N BMAL1 TAD in the presence of 100 µM CBP TAZ2. *, no data due to peak crowding. Open circles, residues that broaden upon addition of TAZ2. e) ^15^N-^1^H HSQC of 100 µM ^15^N BMAL1 TAD L606A/L607A (gray) titrated with CBP TAZ1 (orange to red). f) Differential broadening of 100 µM ^15^N BMAL1 TAD L606A/L607A in the presence of 50 µM CBP TAZ1. g) ^15^N-^1^H HSQC of 100 µM ^15^N BMAL1 TAD L606A/L607A (gray) titrated with CBP TAZ2 (light to dark pink). h) CSPs of 100 µM ^15^N BMAL1 TAD L606A/L607A in the presence of 100 µM CBP TAZ2. *, no data due to peak crowding. Open circles, residues that broaden upon addition of TAZ2.

To determine if the IXXLL motif in the BMAL1 TAD helix remains the key anchoring point for interaction with the TAZ domains, we looked at binding of the ^15^N BMAL1 TAD L606A/L607A mutant in the presence of increasing amounts of CBP TAZ1 (Fig. 6e) or TAZ2 (Fig. 6g). With TAZ1, there are almost no changes to the spectrum of the mutant TAD with one interesting exception: significant broadening of the switch peaks corresponding to just the *cis*-Pro625 isomer (Fig. 6f and inset of spectrum in Fig. 6e). This observation suggests that the TAZ1 domain absolutely requires the IXXLL motif for binding and exhibits preference for the *cis* isomer of the BMAL1 TAD switch. By contrast, the TAZ2 domain retained interaction with the L606A/L607A mutant (Fig. 6g), albeit with lower apparent affinity due to the presence of fast exchange behavior across the TAD, and it did not exhibit an apparent preference for either switch isomer (Fig. 6h).

We were curious if the *cis-*locked TAD would influence its apparent affinity for the TAZ1 domain. To test this, we used fluorescence polarization to assess binding of CBP TAZ1 to a set of [5,6] TAMRA-labeled BMAL1 minimal TAD probes corresponding to the native switch (60% *trans,* 40% *cis*), P625dmP (all *cis*), or P625A (all *trans*), as well as a probe missing all 8 residues of the switch region (Δswitch) (Fig. S5a). Although we noted a modestly lower K_D_ with the *cis*-locked TAD probe (0.54 ± 0.12 µM), it was still similar to the native TAD (0.75 ± 0.14 µM). Truncation of the switch region decreased affinity for TAZ1 by approximately three-fold (2.30 ± 0.23 µM) and reduced the Hill coefficient of binding to 1.1 from approximately to 1.6 with the TADs containing both the helix and switch motifs (Fig. S5b). Collectively, these data demonstrate that the overall affinity of the BMAL1 TAD for multiple domains of CBP/p300 is primarily driven by the IXXLL motif in the BMAL1 TAD helix, supported by recruitment of the switch region.

## Discussion

Changes in CLOCK:BMAL1 activity throughout the day dictate the phase of circadian rhythms^49^, so considerable effort has gone into understanding where the complex acts in the genome^2^, and how it is repressed directly by cryptochromes and other clock proteins^13, 50^ as well as epigenetic regulators^51–53^. However, relatively little is known about the mechanisms of its activation by acetyltransferases such as CBP/p300^19^ and TIP60^54^. Here, we describe how the transactivation domain, or TAD, of BMAL1 interacts with several modular domains in CBP/p300. The potential for multivalent recruitment of CBP/p300 to circadian promoters is increased by binding of CLOCK:BMAL1 to tandem E-box motifs, allowing the presentation of two BMAL1 TADs in close proximity for coactivator binding. We previously demonstrated how CLOCK:BMAL1 binds to a tandem E-box motif in the native *Por* promoter in the context of a nucleosome^5^; here we show that CLOCK:BMAL1 can bind to more internal E-box motifs in the nucleosome if presented in tandem, separated by 6 or 7 base pairs.

Central to regulation of CLOCK:BMAL1 by CBP/p300 are two regulatory motifs in the BMAL1 TAD—the IXXLL motif within the TAD helix and the switch region that comprises the last 8 residues of the protein. Both of these regions in BMAL1 differ in sequence from its paralog BMAL2, which is unable to support circadian rhythms in peripheral organs or cell culture^13, 55^. Moreover, substitutions in these two regions of BMAL1 significantly affect circadian period and amplitude in cell culture^13, 32^, demonstrating the key role they play in mammalian circadian rhythms as the primary transactivation domain of CLOCK:BMAL1.

Our discovery that the two regions of the BMAL1 TAD engage both TF recruitment sites on the KIX domain makes it similar to FOXO3a, which uses two TADs separated by hundreds of amino acids to occupy both sites^28^. However, the two FOXO3a TADs bind both sites on KIX, whereas we saw distinct preferences for the two BMAL1 TAD motifs on the KIX domain. A prior study of BMAL1 TAD interactions with the CBP KIX domain demonstrated that competition for the MLL1 site with a covalent inhibitor led to an approximate 3-fold decrease in affinity^29^, which is similar to the decrease in affinity with deletion of the switch region that we reported earlier^32^. That study^29^ explored only single or double mutations at the c-Myb site, resulting in modest decreases in affinity that minimized the apparent importance of this interface. Based on the significant drop in affinity (∼100-fold) with the loss-of-function triple mutation of the KIX domain c-Myb site or disruption of the IXXLL motif in the BMAL1 TAD helix that we describe here, we propose that these are the primary interfaces for the CBP KIX-BMAL1 TAD interaction.

We found that the BMAL1 TAD helix is also used as the primary interface to bind the CBP TAZ domains. In addition to work showing that the TAD helix is essential for binding other transcriptional regulators^11–14, 56^, this establishes the TAD helix as the central determinant of CLOCK:BMAL1 activity and therefore, transcriptional circadian rhythms in mammals. While we do not yet have structures for these complexes containing tandem-bound CLOCK:BMAL1 and CBP, this work maps out the relevant binding sites and suggests that the landscape of CLOCK:BMAL1 activation is affected by its occupancy of tandem E-box motifs and interactions with multiple domains of CBP/p300 and possibly other transcriptional coactivators^57^. As with the multivalent interactions of p53 and p300^23^, this multivalency might play an important role in helping CLOCK:BMAL1 compete with other TFs to reliably recruit CBP/p300 in cells and generate the transcription-translation feedback that underlies mammalian circadian rhythms.

Although we set out to describe how CLOCK:BMAL1 interacts with its coactivator CBP/p300 in this study, this work has several limitations. First, we only studied three of the known TF-binding domains on CBP. A recent study by Fordyce and colleagues used high-throughput methods to map TF binding to domains of p300 *in vitro*^40^ and identified the same interactions with BMAL1 that we did here in addition to potential interactions with the some of the disordered TF-binding motifs of p300. Additionally, we have not yet directly observed multivalent interactions between CLOCK:BMAL1 and CBP or p300 on DNA or chromatin. However, with new insights into the interactions with CBP/p300 and tandem E-boxes in chromatin provided by this work, it may now be possible to capture these complexes to better understand their central regulatory role within of mammalian circadian rhythms.

## Acknowledgments

We thank Jan Seebacher for assistance with the mass spectrometry data. Support for this work was provided by US National Institutes of Health grants R35 GM141849 (C.L.P.) and R35 GM145255 (S.M.R.), and the Howard Hughes Medical Institute (C.L.P.). N.H.T. funded by the European Research Council under the European Union’s Horizon 2020 research program (NucEM, no. 884331), Novartis Research Foundation, Swiss National Science Foundation (Sinegia-CRS115_186230, SNF 31003A_179541 and SNF 310030_201206) and KFS-4980-02-2020. A.K.M. was supported by a Human Frontier Science Program Long-Term Fellowship, and P.C. by EMBO ALTF 57-2019. D.S. was supported by the California Institute for Regenerative Medicine (CIRM) under award number EDUC4-12759 and the Institute for the Biology of Stem Cells (IBSC) at UC Santa Cruz.

## Author contributions

L.S., A.K.M., N.H.T., and C.L.P. conceived of the study; D.S., L.S., M.R.T., P.C., K.F., N.P., A.K.M., C.L.G., and H.-W.L. performed experiments and analyzed results; M.M. contributed reagents; S.M.R., N.H.T., and C.L.P. oversaw the work and secured funding; C.L.P. wrote the manuscript with assistance from all other authors.

## Materials availability

Materials are available upon request from Carrie Partch or Nicholas Thomä with a materials transfer agreement from their respective institutions.

## Data availability

The electron microscopy (EM) density maps have been deposited in the Electron Microscopy Data Bank (EMD-54868 tandem 6bp, EMD-54869 tandem 7bp). Atomic models were deposited in the RCSB Protein Data Bank (PDB 9SFY tandem 6bp, PDB 9FSZ tandem 7bp). XL-MS data are available through the ProteomeXchange via the PRIDE database with the identifier PRIDE: PXD063793.

## Methods

### Protein expression and purification

#### CLOCK:BMAL1 bHLH-PAS-AB heterodimer

Mouse CLOCK (UniProtKB O08785) bHLH PAS-AB (residues 26–395) and mouse BMAL1 (UniProtKB Q9WTL8-4) bHLH PAS-AB (residues 62–441) were cloned into separate pFastBac vectors as described previously^15^. Briefly, 1–2 L of CLOCK:BMAL1 bHLH-PAS-AB-expressing insect cells (*Spodoptera frugiperda* or Hi5) were pelleted and resuspended in His buffer A (20 mM sodium phosphate buffer pH 8, 200 mM NaCl, 15 mM imidazole, 10% (v/v) glycerol, 0.1% (v/v) Triton X-100 and 5 mM 2-mercaptoethanol). Cells were lysed by cell disruption and subsequent sonication for 3 min (15 sec on, 30 sec off). Lysate was clarified by centrifugation at 45,000 rpm at 4°C for 45 min. Ni-NTA affinity purification was performed on a 5 ml HisTrap FF affinity column (Cytiva). After 15-column washes in His buffer A, the column was further washed with 6.5% His buffer B (20 mM sodium phosphate buffer pH 7.5, 200 mM NaCl, 300 mM imidazole, 10% (v/v) glycerol and 5 mM 2-mercaptoethanol) for 3 column volumes (CV) before being eluted in a 10 CV gradient to 100% His buffer B. The relevant fractions were pooled and cleaved with His-TEV at 4 °C for a minimum of 4 h. The complex was then concentrated to 5–10 ml and re-diluted to 50 ml with heparin buffer A (20 mM sodium phosphate buffer pH 7.5, 50 mM NaCl, 2 mM dithiothreitol and 10% (v/v) glycerol) and loaded onto a HiTrap Heparin HP affinity column (Cytiva). After washing with 5 CV of the above buffer, the column was washed with a further 3 CV of 25% heparin buffer B (20 mM sodium phosphate buffer pH 7.5, 2 M NaCl, 2 mM dithiothreitol and 10% (v/v) glycerol) before eluting with buffer B over an 8 CV gradient. The relevant fractions were purified by Superdex 200 gel filtration chromatography (Cytiva) into 20 mM HEPES buffer pH 7.5, 125 mM NaCl, 5% (v/v) glycerol and 2 mM TCEP Small aliquots of the complex were quick frozen in liquid nitrogen and stored at -70°C.

#### BMAL1 TAD

Mouse BMAL1 (UniProtKB Q9WTL8-4) TAD (residues 579-626) was cloned into a bacterial expression plasmid based on the pET22b vector backbone from the parallel vector series^58^. The L606A/L607A mutation was generated previously^13^. All TAD constructs possessed an N-terminal TEV cleavable His6 NusA extra-linker (HNXL) solubilizing tag and ampicillin resistance. Rosetta (DE3) *E. coli* harboring TAD plasmids were grown to an OD600 of ∼0.6-0.9 in the presence of ampicillin (100 µg/mL) and chloramphenicol (35 µg/mL). Protein expression was induced with 0.5 mM isopropyl β-D-1-thiogalactopyranoside (IPTG) and allowed to proceed for 16-18 hours at 18°C in either Luria Broth (LB) or M9 minimal medium containing 1 g/L ^15^NH_4_Cl to generate uniformly ^15^N-labeled proteins for NMR spectroscopy. Cells were lysed in with an Emulsiflex C-3 cell disruptor (Avestin) in Buffer A containing 50 mM Tris pH 7.5, 300 mM NaCl and 20 mM imidazole. The soluble fraction of *E. coli* lysates was passed over Ni-NTA resin (QIAGEN), washed thoroughly, and eluted using 250 mM imidazole. Fractions of interest were buffer exchanged into lysis buffer using a stirred-cell pressure concentrator with an Amicon 3 kDa molecular weight cutoff (MWCO) filter (Merck Millipore). Proteolysis was performed with His_6_-tagged TEV protease overnight at 4°C and cleaved protein was retained from the flow-through of a Ni-NTA column. The protein was further purified on a preparative grade Superdex 75 16/600 size-exclusion column (Cytiva) pre-equilibrated with NMR buffer (10 mM MES pH 6.5 and 50 mM NaCl). Small aliquots of the protein were quick frozen in liquid nitrogen and stored at -70°C.

#### CBP KIX

The bacterial expression plasmid encoding the mouse CBP (UniProt P45481) KIX (residues 586-672) was a gift from Peter Wright (Scripps Research Institute). The Y650A/A654Q/Y658A (3mut) mutation introduced by site-directed mutagenesis and verified by sequencing. Rosetta (DE3) *E. coli* cells harboring CBP KIX plasmids were grown as the BMAL1 TAD above and purification proceeded similarly, except that CBP KIX has native histidine residues that allow for the purification of the tagless protein using Ni-NTA resin.

#### CBP TAZ1

Mouse CBP (UniProt P45481) TAZ1 (residues 340-439) was cloned into the pET22b vector without a tag and with ampicillin resistance. Rosetta (DE3) *E. coli* were grown to an OD600 of ∼0.6-0.9 in the presence of ampicillin (100 µg/mL) and chloramphenicol (35 µg/mL) at 37°C, and then expression was induced with 1 mM IPTG and 50 µM ZnSO_4_ and grown for 4 hours at 37°C. Bacterial pellets were collected by centrifugation, and the cells were resuspended in Buffer A (50 mM Tris pH 7.5, 300 mM NaCl, 20 mM imidazole, 20 mM DTT, 50 µM ZnSO_4_) and lysed by high pressure on ice. The crude extract was separated by centrifugation at 19,000 rpm at 4°C. The insoluble pellet containing TAZ1 protein was washed once with Buffer A and repelleted through centrifugation. The final insoluble pellet was resuspended in Buffer B (10 mM Tris pH 7.5, 20 mM NaCl, 6 M Urea, 20 mM DTT, 50 µM ZnSO_4_). Complete resuspension of the inclusion bodies was achieved through sonication (40% amplitude, 15 sec on, 30 sec off) and vortexing, and the solubilized pellet was centrifuged at 19,000 rpm for 30 mins to remove insoluble particulates. The supernatant was filtered with a 0.40-micron membrane and passed over a SP sepharose cation exchange column (Cytiva). Protein was eluted with a gradient from Buffer C (50 mM Tris pH 7.5, 20 mM NaCl, 20 mM DTT, 50 µM ZnSO_4_) to Buffer D (50 mM Tris pH 7.5, 1 M NaCl, 20 mM DTT, 50 µM ZnSO_4_). After elution, three equivalents of ZnSO_4_ were added to fractions containing TAZ1 to properly refold the denatured protein. Refolded protein was then run on a preparative Superdex 75 16/600 size-exclusion column (Cytiva) in NMR Buffer (10 mM MES pH 6.5, 50 mM NaCl, 2 mM DTT). Small aliquots of the protein were quick frozen in liquid nitrogen and stored at -70°C.

#### CBP TAZ2

Mouse CBP (UniProt P45481) TAZ2 (residues 1764-1855) was expressed in *E. coli* (BL21) cells as an MBP fusion protein and purified using amylose sepharose affinity chromatography and S-sepharose ion exchange chromatography as described in^59^. TAZ2 contained cysteine to alanine point mutations (C1776A, C1784A, C1827A, C1828A) that increase solubility and stability^60^. ZnSO_4_ was kept in the media for TAZ2 expression at a concentration of 1 μM. TAZ2 was further purified with Superdex 75 size exclusion chromatography (Cytiva) and buffer exchanged into NMR buffer (10 mM MES pH 6.5, 50 mM NaCl). Small aliquots of the protein were quick frozen in liquid nitrogen and stored at -70°C.

### Peptide synthesis

The linker-switch peptide (residues 613-626 of mouse BMAL1) was synthesized by Pierce and provided at 95% purity. The minimal TAD peptides (residues 594-626) for WT, P625dmP or P625A, and the Δswitch variant (594-F619Y with an amidated C-terminus) were synthesized by Bio-Synthesis with a 5,6-TAMRA fluorescent probe covalently attached to the N-terminus and provided at 95% purity. 8-mer switch peptides FSDLPWPL and the *trans-*locked FSDLPWAL were synthesized by solid phase peptide synthesis on 3-chlortrityl resin with standard Fmoc chemistry. One or two 1:4:4:4:6 molar ratio of Resin:HBTU:HOAT:Fmoc-AA-OH:DiPEA coupling reactions were performed in DMF for each amino acid addition. The *cis*-locked peptide FSDLPWdmPL (dmP, 5,5- dimethyl L-proline) was synthesized by solid phase peptide synthesis using standard Fmoc chemistry. Coupling of dmP onto the Leu-Resin was performed using 1:2:2:4 molar ratio of Resin:HATU:Fmoc-dmP-OH:DiPEA, and coupling of the Trp onto the Resin-Leu-dmP was performed using 1:3.8:4:6 molar ratio of Resin:COMU:Fmoc-Trp-Boc-OH:DiPEA; all other coupling reactions were performed using HBTU/HOAT as described above. Naturally occurring Fmoc-protected amino acids were purchased from Fluka, Nova Biochem, AAPPTec, or Sigma Aldrich. Fmoc-dmP was purchased from PolyPeptide Group (cat. # FA21702). Peptides were purified by reverse phase C18 HPLC; purity (>90%) and identity were verified by MS/MS on a Waters HPLC-MS/MS system.

### Nucleosome assembly and complex formation

#### Human octamer histone expression, purification, and reconstitution

Human histones were expressed and purified as described previously^61^. Lyophilized histones were mixed at equimolar ratios in 20 mM Tris-HCl pH 7.5 buffer containing 7 M guanidine hydrochloride and 20 mM 2-mercaptoethanol. Samples were dialyzed against 10 mM Tris-HCl pH 7.5 buffer containing 2 M NaCl, 1 mM EDTA, and 2 mM 2-mercaptoethanol. The resulting histone complexes were purified by Superdex 200 size exclusion chromatography (Cytiva).

#### DNA preparation

DNA for medium to large-scale individual nucleosome purifications was generated by Phusion (Thermo Fisher Scientific) PCR amplification. The resulting DNA fragment was purified by a MonoQ column (Cytiva). All purified DNA was concentrated and stored at - 20°C in 20 mM Tris-HCl pH 7.5 until use.

#### Nucleosome assembly

The DNA and the histone octamer complex were mixed in a 1:1.5 molar ratio in the presence of 2 M KCl. The samples were dialyzed against refolding buffer (RB) high (10 mM Tris-HCl pH 7.5, 2 M KCl, 1 mM EDTA, and 1 mM DTT). The KCl concentration was gradually reduced from 2 M to 0.25 M using a peristaltic pump with RB low (10 mM Tris-HCl pH 7.5, 250 mM KCl, 1 mM EDTA, and 1 mM DTT) at 4°C. The reconstituted nucleosomes were incubated at 55°C for 2 h followed by purification on a MonoQ 5/50 ion exchange column (Cytiva) and dialyzed into 20 mM Tris-HCl pH 7.5 and 500 μM TCEP overnight. Nucleosomes were concentrated and stored at 4°C.

### Cryo-electron microscopy

#### Cryo-EM sample preparation

Nucleosomes were mixed with CLOCK:BMAL1 (1:3) in ∼100 μL volume and incubated at RT for 30 min in a binding buffer containing 20 mM HEPES pH 7.4, 1 mM MgCl_2_, 10 mM KCl, and 0.5 mM TCEP. The sample was then subjected to cross-linking using the GraFix method^62^. For GraFix cross-linking, the CLOCK:BMAL1-NCP complexes were layered on top of a 10%–30% (w/v) sucrose gradient (20 mM HEPES pH 7.4, 50 mM NaCl, 1 mM MgCl_2_, 10 mM KCl, 0.5 mM TCEP) with an increasing concentration (0-0.34% w/v) of glutaraldehyde (EMS) and subjected to ultracentrifugation (Beckman SW40Ti rotor, 30000 rpm, 18 h, 4°C). After centrifugation, 100 μL fractions were collected from the top of the gradient and peak fractions were analyzed by native PAGE. The peak fractions were combined, and sucrose was removed by dialysis into GraFix buffer (20 mM HEPES pH 7.4, 50 mM NaCl, 1 mM MgCl_2_, 10 mM KCl, 0.5 mM TCEP). The resulting sample was concentrated with an Amicon Ultra 0.5 mL 30kDa MWCO centrifugal filter (Merck Millipore) to ∼5-7 μM nucleosomes as determined by measuring the DNA concentration at absorbance at 260 nm. After concentration, 3.5 µL of sample was applied to glow discharged Quantifoil holey carbon grids (R 1.2/1.3 200-mesh, Quantifoil Micro Tools). Glow discharging was carried out in a Solarus plasma cleaner (Gatan) for 15 s in an H_2_/O_2_ environment. Grids were blotted for 3 s at 4°C at 100% humidity in a Vitrobot Mark IV (FEI), and then immediately plunged into liquid ethane.

#### Cryo-EM data collection

Data were collected automatically with EPU 3 (Thermo Fisher Scientific) on a Cs-corrected (CEOS GmbH) Titan Krios (Thermo Fisher Scientific) electron microscope operated at 300 kV. The acquisition was performed at a nominal magnification of 75,000 – 96,000× with a Falcon 4 direct electron detector (Thermo Fisher Scientific). All datasets were recorded with an accumulated total dose of 50 e–/Å^2^ and the exposures were fractionated into 50 frames. The targeted defocus values ranged from -0.25 to -2.5 μm.

#### Cryo-EM image processing

Real-time evaluation along with acquisition with EPU 3 (Thermo Fisher Scientific) was performed with CryoFLARE 1.10^63^,. CryoFLARE utilizes the Relion 3 motioncorr implementation^64^ and gCTF^65^ for generation of dose weighted and motion corrected averages and their corresponding CTF fits. All datasets were further processed in cryoSPARC v3^66^ and partially also in Relion 3.0^64^. In detail, in the case of CLOCK:BMAL1-NCP^tandem^ ^6bp^, 2 rounds of 2D classification were performed in cryoSPARC, followed by *ab initio* reconstruction and heterogenous refinement. The selected particles were subdivided into 5 classes by another round of heterogenous refinement and the selected class was further refined by non-uniform refinement. The final particle subset was obtained by performing two rounds of 3D variability analysis using particle clustering, with a mask covering the full complex. The final map was then obtained by non-uniform refinement of these particles.

In the case of CLOCK:BMAL1-NCP^tandem^ ^7bp^, 2 rounds of 2D classification in cryoSPARC, followed by *ab initio* reconstruction, heterogenous refinement as well as a round of non-uniform refinement were performed before the particles were imported into Relion. There, the particles were refined with a reference map from cryoSPARC before being classified into 15 classes by 3D classification. Two more rounds of refinements were perfomed on the selected particles and then they were transferred to cryoSPARC for a 3D variability analysis with particle clustering, followed by a final round of non-uniform refinement. The resolution values reported for all reconstructions are based on the gold-standard Fourier shell correlation curve (FSC) at 0.143 criterion^67, 68^ and all the related FSC curves are corrected for the effects of soft masks using high-resolution noise substitution^69^.

#### Model building and refinement

In the case of the internal tandem CLOCK:BMAL1 structure where the two heterodimers are 7 bp apart, the NCP template (PDB 6T93)^25^ was fitted into the density and the DNA sequence was adjusted. The two heterodimers (without the nucleosome) from the *Por* promoter structure^5^ were fitted into the density for the two bHLH domains. As the density of the PAS domains of both heterodimers was ambiguous, they were both deleted and the nucleosome, DNA and bHLH domains were refined in ISOLDE^70^ and further refined with PHENIX^71^.

In the case of the internal tandem CLOCK:BMAL1 structure where the two heterodimers are 6 bp apart, the previous model was used and the DNA sequence was adjusted. The internal CLOCK:BMAL1 dimer was then deleted and replaced by a CLOCK:BMAL1 template (fusion of DNA/bHLH domains (PDB 4H10) and PAS domains (PDB 4F3L)) by aligning the E-box oligo on the E-box within the nucleosomal DNA. The DNA oligo was then deleted and the bHLH domains and PAS domains were fitted into the density. The orientation of the bHLH domain and the position of the PAS domains were guided by the crosslinks obtained from cross-linking mass spectrometry. The PAS domains of the external heterodimer were not modelled due to limited resolution. Finally, the model was refined using adaptive distance restraints in ISOLDE^70^ and further refined with PHENIX^71^ using coordinate restraints.

### Crosslinking/mass spectrometry

#### Sample preparation

CLOCK:BMAL1 and the nucleosomes were mixed in a 1.5:1 ratio in MS sample buffer (50 mM HEPES pH 7.5, 150 mM NaCl, 500 μM TCEP) and incubated at RT for ∼1 h. In the meantime, an aliquot of DSSO XL reagent (Disuccinimidyl sulfoxide, Thermo Fisher Scientific, cat. #A33545) was warmed up to RT and diluted to a 100 mM stock concentration in anhydrous DMSO by shaking for 5 min, 400 rpm. After incubation, the sample was transferred to an Amicon Ultra 10kDa MWCO concentrator (Merck Millipore), DSSO was added and the cross-linking reaction mix was incubated for 1 h at 10°C, while shaking at 400 rpm. The excess crosslinker was quenched by adding 1 M Tris pH 6.8 (50 mM final concentration) and incubating for an additional hour at RT, 400 rpm. The sample was centrifuged (5 min, 14,000 g) to remove XL reagent and 400 µL of fresh 8 M urea in 50 mM HEPES, pH 8.5 for denaturing and washing were added. This step was repeated 2x. Next, reduction/alkylation buffer (50 mM TCEP, 100 mM 2-chloroacetamide) was added (5 mM and 10 mM final concentration respectively) and the sample was incubated for 30 min while shaking at 400 rpm. It was centrifuged for 5 min at 14,000 g and 400 µL of fresh 8 M urea were added for denaturing and washing. The sample was centrifuged again for 5 min at 14,000 g. This step was repeated 2x with a final centrifugation step of 15 min instead of 5 min to concentrate the sample to ∼30 µL. Lys-C was added (0.2 µg/µL stock, 1:100 enzyme to protein ratio) and the sample was digested for 1.5 h at RT while shaking. The sample was diluted 4-fold with 50 mM HEPES, pH 8.5. Then, trypsin (0.2 mg/mL stock, 1:100 enzyme to protein ratio) was added and the sample was incubated overnight at 37°C, while shaking at 400 rpm. An additional aliquot of trypsin and acetonitrile to a final concentration of 5% was added the next day and the sample was incubated for another 4 h at 37°C, while shaking at 400 rpm. The sample was transferred into an Eppendorf tube, TFA was added (1% final concentration) and the sample was briefly sonicated and spun down for 5 min at 20,000 g. The supernatant was desalted using a PreOmics iST-NHS kit and concentrated in a speedvac. Samples were reconstituted with 0.1% TFA in 2% acetonitrile.

#### Sample analysis

Samples were analyzed by LC-MS in two ways:

1. The equivalent of ca. 1 µg peptides per sample were loaded onto a uPAC C18 trapping column and then separated on a 50 cm uPAC C18 HPLC column (connected to an EASY Spray source (all Thermo Fisher Scientific, columns formerly from Pharmafluidics) connected to an Orbitrap Fusion Lumos. The following chromatography method was used: 0.1% formic acid (buffer A), 0.1% formic acid in acetonitrile (buffer B), flow rate 500 nl/min, gradient 240 min in total, (mobile phase compositions in % B): 0–5 min 3–7%, 5–195 min 7–22%, 195–225 min 22–80%, 225–240 min 80%.
2. The equivalent of ca. 5 µg peptides per sample were loaded onto a Vanquish Neo chromatography system with two-column setup. Samples were injected with 1% TFA and 2% acetonitrile in H_2_O onto a trapping column at a constant pressure of 1000 bar. Peptides were chromatographically separated at a flow rate of 500 nl/min using a 3 h method, with a linear gradient of 2-9% B in 5 min, followed by 9-28% B in 120 min, followed by 28-100% B in 20min, and finally washing for 15 min at 100% B (Buffer A: 0.1% formic acid; buffer B: 0.1 formic acid in 80% acetonitrile) on a 15 cm EASY Spray Neo C18 HPLC column mounted on an EASY Spray source connected to an Orbitrap Eclipse mass spectrometer with FAIMS (Thermo Fisher Scientific).

In either case, the mass Lumos or Eclipse spectrometer was operated in “MS2_MS3” mode, essentially according to prior methods^72^. On the Orbitrap Fusion Lumos mass spectrometer, peptide MS1 precursor ions were measured in the Orbitrap at 120k resolution. On the Orbitrap Eclipse, three experiments were defined in the MS method, with three different FAIMS compensation voltages, -50, -60 and -75 V, respectively, to increase the chances for more highly charged peptides, i.e. cross-linked peptides, to be identified.

For each experiment, peptide MS1 precursor ions were measured in the Orbitrap at 60k resolution. In either case, the MS’ Advanced peak determination (APD) feature was enabled, and those peptides with assigned charge states between 3 and 8 were subjected to CID–MS2 fragmentation (25% CID collision energy), and fragments detected in the Orbitrap at 30k resolution. Data-dependent HCD-MS3 scans were performed if a unique mass difference (Δm) of 31.9721 Da was found in the CID–MS2 scans with detection in the ion trap (35% HCD collision energy).

MS raw data were analyzed in Proteome Discoverer version 2.5 (Thermo Fisher Scientific) using a Sequest^73^ database search for linear peptides, including crosslinker modifications, and an XlinkX^72^ search to identify cross-linked peptides. MS2 fragment ion spectra not indicative of the DSSO crosslink delta mass were searched with the Sequest search engine against a custom protein database containing the expected protein components, as well as a database built of contaminants commonly identified during in-house analyses, from MaxQuant^74^, and cRAP (ftp://ftp.thegpm.org/fasta/cRAP), using the target-decoy search strategy^75^. The following variable crosslinker modifications were considered: DSSO Hydrolysed/+176.014 Da (K); DSSO Tris/+279.078 Da (K), DSSO alkene fragment/+54.011 Da (K); DSSO sulfenic acid fragment/+103.993 Da (K), as well as Oxidation/+15.995 Da (M). Carbamidomethyl/+57.021 Da (C) was set as a static modification. Trypsin was selected as the cleavage reagent, allowing a maximum of two missed cleavage sites, peptide lengths between 4 or 6 and 150, 10 ppm precursor mass tolerance, and 0.02 Da fragment mass tolerance. PSM validation was performed using the Percolator node in PD and a target FDR of 1%.

XlinkX version 2.0 was used to perform a database against a custom protein database containing the expected complex components to identify DSSO-cross-linked peptides and the following variable modification: DSSO Hydrolyzed/+176.014 Da (K); Oxidation/+15.995 Da (M). Crosslink-to-spectrum matches (CSMs) were accepted above an XlinkX score of 40. Crosslinks were grouped by sequences and link positions and exported to xiNET^76^ format to generate cross-link network maps. The mass spectrometry proteomics data have been deposited to the ProteomeXchange Consortium via the PRIDE partner repository with the dataset identifier PXD063793.

#### Mapping crosslinks onto structural models

Crosslinks were mapped to the structural models with an in-house script for PyMOL and the ChimeraX plugin XMAS^77^. Xwalk was used to calculate solvent accessible surface distances^78^.

### Mass photometry

For measuring nucleosomes or nucleosome complexes, microscope coverslips were treated with 10 µL of poly-L-lysine for 30 s, rinsed with Milli-Q and dried under an air stream. Prior to mass photometry measurements, protein dilutions were made in MP buffer (20 mM Tris-HCl pH 7.5, 100 mM KCl, 0.5 mM TCEP) and nucleosome:CLOCK:BMAL1 complexes were mixed in a 1:6 ratio and incubated for 30 min at room temperature. Data was acquired on a Refeyn OneMP mass photometer. First, 18 µL of MP buffer was introduced into the flow chamber and focus was determined. Then 2 µL of protein solution were added to the chamber and movies of 60 s or 90 s were recorded. Nucleosomes (NCP^tandem^ ^6bp^ and NCP^tandem^ ^7bp^) and CLOCK:BMAL1 bHLH-PAS-AB were measured individually at 20 nM (final concentration) and then in complex at 10 nM and 60 nM, respectively. Each sample was measured at least two times independently. All acquired movies were processed and molecular masses were analyzed using Discover 2.3, based on a standard curve created with BSA and Thyroglobulin.

### NMR spectroscopy

NMR experiments were conducted on a Varian INOVA 600-MHz spectrometer equipped with ^1^H, ^13^C, ^15^N triple resonance, Z-axis pulsed field gradient cryoprobe. All NMR data were collected at 25°C. NMR data were processed using NMRPipe/NMRDraw^79^. ^15^N-^1^H HSQC spectra CBP KIX were performed in 330 μL volume in a Shigemi tube with 100 μM ^15^N CBP KIX in NMR buffer (10 mM MES pH 6.5, 50 mM NaCl) and 10% (v/v) D_2_O in the presence of stepwise additions of concentrated stocks of BMAL1 TAD or truncated peptides. ^15^N-^1^H HSQC spectra of BMAL1 TAD WT or L606A/L607A were performed in 330 μL volume in a Shigemi tube with 100 μM ^15^N BMAL1 TAD in NMR buffer (10 mM MES pH 6.5, 50 mM NaCl) and 10% (v/v) D_2_O in the presence of stepwise additions of CBP KIX (WT or 3mut), TAZ1, or TAZ2, followed by concentration in an Amicon Ultra centrifugal filter with a 3 kDa MWCO (Merck Millipore). Chemical shift assignments of BMAL1 TAD (entry 25280) and CBP KIX (entry 6750) were obtained from the Biological Magnetic Resonance Data Bank (BMRB); CBP KIX assignments at pH 5.5 were transferred to the spectrum at pH 6.5 via a 3-step pH titration. Titration data for protein interactions were analyzed with NMRViewJ (One Moon Scientific) using chemical shift perturbations defined by the equation Δδ_TOT_ = [(Δδ^1^H)^2^ + (χ(Δδ^15^N)^2^]^½^ and normalized with the scaling factor χ = 0.5^80^.

### Isothermal titration calorimetry

ITC measurements were obtained as previously described^13^. Briefly, proteins were extensively dialyzed at 4°C in 10 mM MES pH 6.5, 50 mM NaCl using a 2 kDa molecular weight cutoff filter dialysis tubing (Spectrum Labs) prior to collecting ITC data. ITC was performed on a MicroCal VP-ITC calorimeter at 25°C with a stir speed of 177 rpm, reference power of 10 μCal/sec and 10 μL injection sizes. Protein ratios for the cell and syringe for the ITC assays (3 independent runs for each complex) were 220-230 μM CBP KIX WT or 3mut titrated into 20-25 μM BMAL1 TAD WT or L606A/L607A as indicated (stoichiometry (n) = 0.8-1.1). All data were best fit by a one-site binding model using Origin software.

### Fluorescence polarization

The BMAL1 minimal TAD WT, P625A, P625dmP (594-626) and Δswitch (594-F619Y) probes were purchased from Bio-Synthesis Inc. with a [5,6] TAMRA fluorescent probe covalently attached to the N-terminus; the C-terminus of the Δswitch peptide was amidated, while the others were left as a free carboxyl group. Equilibrium binding assays (Fig. 5b) with CBP KIX, TAZ1, and TAZ2 and competition assay with CBP KIX:TAD WT titrated with KG-501 (Fig. 5i) were performed in 10 mM MES pH 6.5, 50 mM NaCl and 0.05% (v/v) Tween-20. Concentrated stocks of BMAL1 TAD probes were stored between 15-200 μM at -70°C and diluted in assay buffer to 50 nM alone (binding assays) or to 10 nM bound to 2 µM CBP KIX (competition assay) and in the presence of increasing concentrations of KG-501. Plates were incubated at room temperature for 10-20 minutes prior to analysis. Binding was monitored by changes in fluorescence polarization with an EnVision 2103 multilabel plate reader (Perkin Elmer) with excitation at 531 nm and emission at 595 nm. The Hill coefficient and equilibrium binding dissociation constant (K_D_) were calculated by fitting the dose-dependent change in millipolarization (Δmp) to a one-site specific binding model (binding assays) or an [inhibitor] vs. response (three parameters) model (competition assay) in Prism (GraphPad). Data shown for equilibrium binding assays are from a representative experiment (n=4 replicates) of three independent assays (Fig. 5b). Data shown for competition assay are from a representative experiment (n = 2 replicates) of two independent assays (Fig. 5i). The [5,6] TAMRA labeled minimal BMAL1 TAD bound to CBP KIX in the absence of KG-501 (protein complex alone) producing a fluorescence signal of ∼ 100 mp and a K_D_ = 2 μM was determined by titration of CBP KIX. Because KG-501 was solubilized in DMSO, a 2% DMSO control (highest concentration used in KG-501 titration) was performed and it induced a ∼10% mp decrease in FP signal relative to the complex alone. The IC_50_ reported is the mean of two independent assays and the K_i_ was calculated using the Cheng-Pursoff equation setting the K_D_ = 2 μM for the BMAL1 TAD-CBP KIX interaction.

### Cell culture and assays

HEK293T cells were purchased from ATCC (cat. #CRL-3216) were cultured in 10% DMEM (i.e. 10% (v/v) FBS) and 1X penicillin-streptomycin (Thermo Fisher) at 37°C in an incubator humidified with 5% CO_2_. Forskolin (CAS 66575-29-9, cat. #F3917, Sigma Aldrich), KG-501/naphthol-AS-E-phosphate (CAS 18228-17-6, cat. #70485, Sigma Aldrich), XX-650-23 (CAS 117739-40-9, cat. #AOB31725, Aobious), and 666-15 (CAS 1433286-70-4, cat. #AMBH97F07B56, Sigma Aldrich) were stored as recommended by the manufacturer and assessed by 1D NMR before use.

For *Per1*-luc reporter assays with full-length CBP and domain deletions, Plasmids expressing CBP with specific domain deletions generated from pcDNA4B Flag CBP WT. The NEBuilder assembly tool was used to design primers that would generate the desired deletion constructs. NEBuilder HiFi DNA assembly was then performed as per the manufacturer’s protocol. Formation of the desired plasmid was confirmed by restriction digest, sanger sequencing, and whole plasmid sequencing for all constructs.

48-well plates with HEK293T cells were subsequently transfected with LT-1 (Mirus) in quadruplicate as follows: for *Per1-*luc alone, 5 ng pGL3 *Per1*-luciferase^8^ and 600 ng empty pcDNA4 vector; for CLOCK:BMAL1 activity, 5 ng pGL3 *Per1*-luciferase, 100 ng pSG5 Clock, 100 ng pSG5 Bmal1, and 400 ng empty pcDNA4 vector. The remaining wells contained 5 ng pGL3 *Per1*-luciferase, 100 ng pSG5 Clock, 100 ng pSG5 Bmal1 and the indicated quantity of CP WT or domain deletion plasmid, with the total DNA concentration kept constant using empty pcDNA4 vector. Cells were allowed to grow for 48 hours before imaging with an ALLIGATOR (Cairn Research Ltd). Luciferase expression was determined using the FIJI image processing package.

For *Per1-*luc reporter assays with inhibitors, 48-well plates with HEK293T cells were transfected with LT-1 (Mirus) and the following plasmids in duplicate: 5 ng pGL3 *Per1*-luciferase^8^ and 395 ng empty pcDNA4 vector; for CLOCK:BMAL1 activity, 5 ng pGL3 *Per1*-luciferase, 100 ng pSG5 Clock, 100 ng pSG5 Bmal1, and 195 ng empty pcDNA4 vector. Cells were treated with compounds as indicated or 1% DMSO 24 hours after transfection for 16 hours prior to harvesting. For forskolin treatment, 10 μM forskolin was added 2 hours prior to harvesting. Cells were lysed in Passive Lysis Buffer (New England Biolabs) according to the protocol and luciferase activity was measured with Bright-Glo reagent (New England Biolabs) on an EnVision plate reader (Perkin Elmer). Each reporter assay was repeated at least three times.

### Western blotting

HEK293T cells were cultured as above and plated in 6-well plates. Cells were transfected with LT-1 (Mirus) in a 3 μL:1 μg LT-1:DNA ratio with 1 μg of pcDNA 4B Flag CBP WT or deletion mutant plasmid/well and allowed to grow 48-72 hours after transfection. Cells were transferred into an Eppendorf tube with PBS, pelleted, and lysed for 10 minutes in 100 μL of lysis buffer (20 mM Tris pH 7.5, 150 mM NaCl, 1 mM TCEP, 0.1% NP-40 and EDTA-free protease inhibitors (Roche). Following 30 minutes centrifugation at 19000 RPM at 4°C, the supernatant added to 6x Laemmli buffer. A total of 19 μL of extracts/well were run on 7.5% SDS-PAGE, transferred onto a nitrocellulose membrane and blocked for 1 hour at room temperature in TBST with 5% milk. Western blotting was performed using the following primary antibodies: the DYKDDDDK Tag antibody, HRP-conjugated (GenScript cat. # A01428-100, lot # 2105K001) and GAPDH antibody (G-9) (Santa Cruz Biotech cat. # sc-365062) and the following secondary antibody: anti-mouse IgG-HRP (Cell Signaling cat. # 58802S). Immobilon Western Chemiluminescent HRP substrate (Merck) was used for chemiluminescent detection.

### Statistical analysis

Where applicable, statistical parameters including sample size, precision measures (standard deviation, s.d.) and statistical significance are reported in the Figures and corresponding Figure Legends, calculated in Prism (GraphPad).

