## Supplemental Materials for "Tandem association of CLOCK:BMAL1 complexes on DNA enables recruitment of CBP/p300 through multivalent interactions"

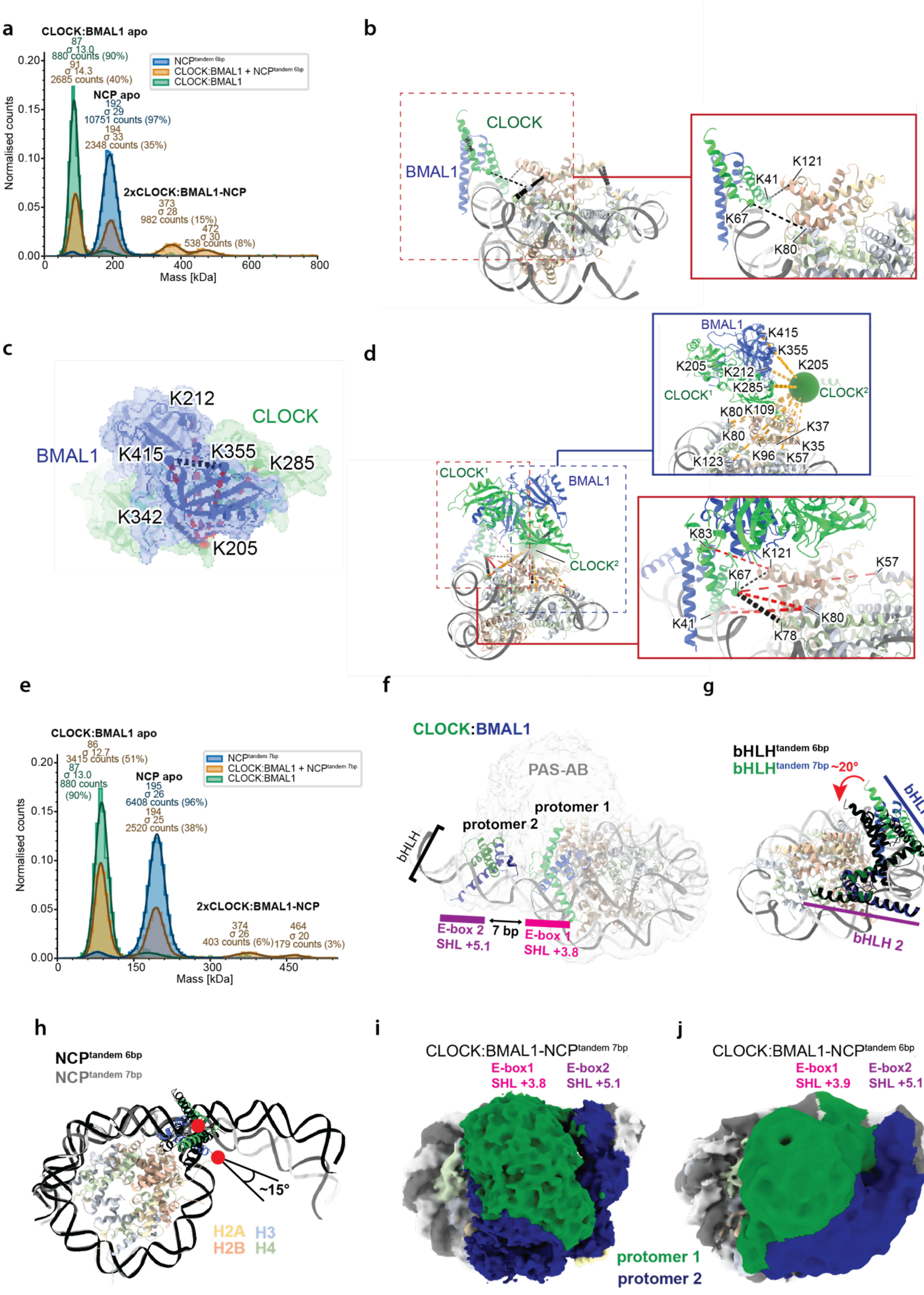
a
CLOCK:BMAL1 apo
NCP apo
2xCLOCK:BMAL1-NCP
Normalised counts
Mass [kDa]
b
CLOCK
BMAL1
K121
K41
K67
K80
c
K212
CLOCK
BMAL1
K415
K355
K285
K342
K205
d
CLOCK¹
CLOCK²
BMAL1
e
CLOCK:BMAL1 apo
NCP apo
2xCLOCK:BMAL1-NCP
f
CLOCK:BMAL1
PAS-AB
protomer 1
protomer 2
bHLH
E-box 2
SHL +5.1
7 bp
E-box 1
SHL +3.8
g
bHLH 1
bHLH 2
~20°
h
~15°
H2A
H3
H2B
H4
i
CLOCK:BMAL1-NCPtandem 7bp
E-box1
SHL +3.8
E-box2
SHL +5.1
protomer 1
protomer 2
j
CLOCK:BMAL1-NCPtandem 6bp
E-box1
SHL +3.9
E-box2
SHL +5.1

**Figure S1: Characterization of CLOCK:BMAL1-NCP complexes.** a) Molecular mass distribution histogram of CLOCK:BMAL1-NCP$^{tandem\ 6bp}$. CLOCK:BMAL1 and the nucleosomes were first measured individually at 20 nM and in a 1:6 ratio. CLOCK:BMAL1 primarily forms complexes with nucleosomes in a 1:2 complex with a minority species of 1:3 stoichiometry. b) Cross-links between the bHLH domain of CLOCK:BMAL1 and histones allow the specific identification of CLOCK and BMAL1. c) Cross-links mapped to the PAS domains of a single CLOCK:BMAL1 heterodimer. Sterically incompatible cross-links are depicted in red. d) Tentative model for the first CLOCK:BMAL1 protomer. CLOCK K205 from the second protomer, which makes multiple cross-links is depicted as a sphere (blue box). Cross-links did not allow unambiguous model building and suggest that multiple conformations for the PAS domains of the second protomer are possible. Putative inter-CLOCK:BMAL1 cross-links would be sterically incompatible when mapped to a single heterodimer (see panel c). e) Molecular mass distribution histogram of CLOCK:BMAL1-NCP$^{tandem\ 7bp}$. CLOCK:BMAL1 and the nucleosomes were first measured individually at 20 nM and in a 1:6 ratio. CLOCK:BMAL1 primarily forms complexes with nucleosomes in a 1:2 complex with a minority species of 1:3 stoichiometry. f) Map-model overlay for CLOCK:BMAL1-NCP$^{tandem\ 7bp}$. The resolution of the map only allowed confidently building of the bHLH domains of both protomers. g) Comparison of the two models, aligned on the second bHLH domain at SHL+5.1 indicates a rotation of ~ 20º of the first bHLH domain. h) Comparison of the DNA trajectory between CLOCK:BMAL1-NCP$^{tandem\ 6bp}$ and CLOCK:BMAL1-NCP$^{tandem\ 7bp}$. Only the internal bHLH domain is depicted for clarity. i-j) Top view of the cryo-EM map obtained for CLOCK:BMAL1 binding to tandem E-boxes with 7 bp spacing (i) or 6 bp spacing (j).

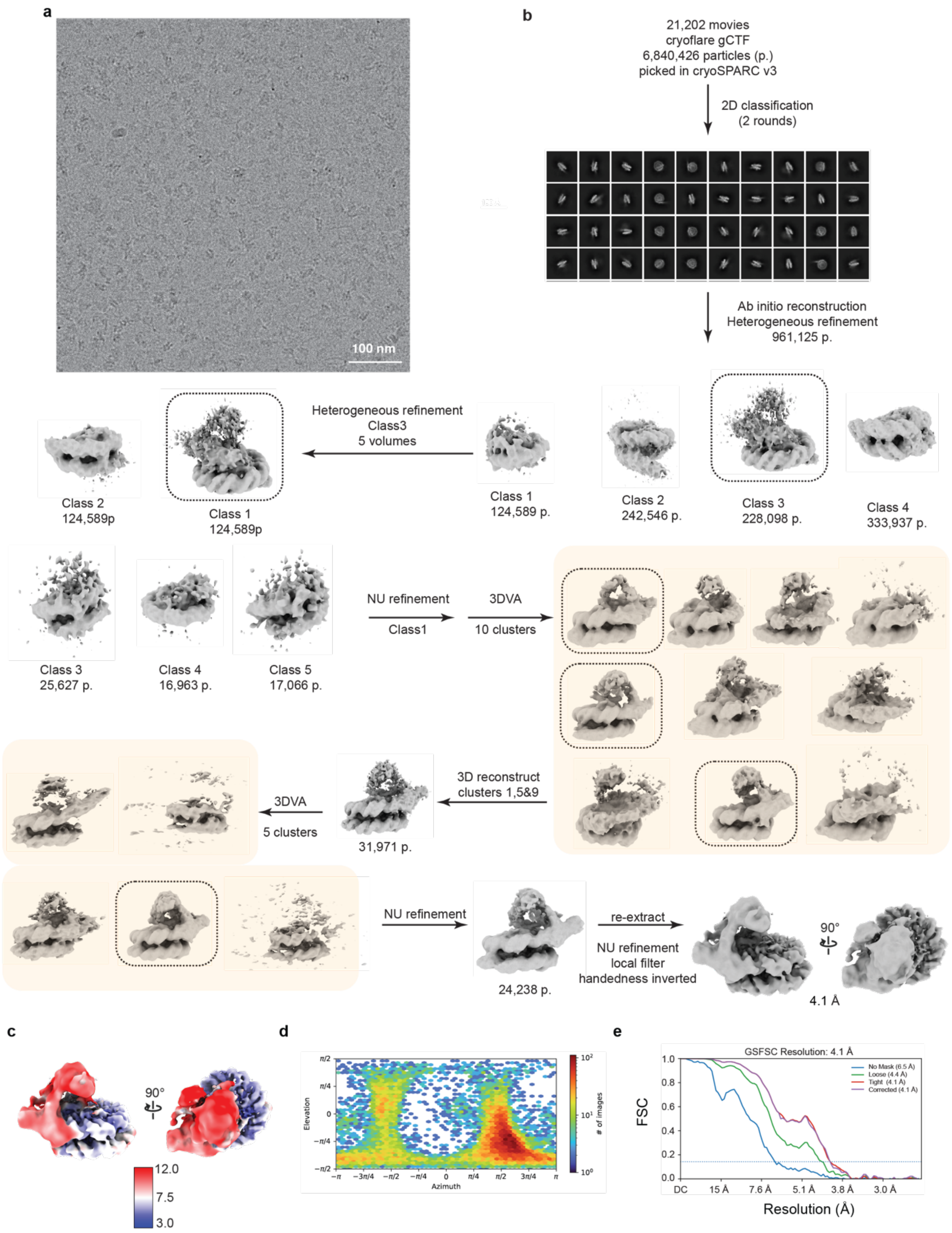

**Figure S2: cryo-EM data processing workflow for CLOCK:BMAL1-NCP$^{\text{tandem 6bp}}$.** a) Representative cryo-EM micrograph. b) See methods. The movies were pre-processed within cryoFLARE gCFT and the resulting micrographs were imported in cryoSPARC v3 for further processing. Multiple rounds of classification, refinement and 3D variability analysis followed by non-uniform refinement (NU refinement) resulted in the final map. The dashed boxes indicate the maps and sets of particles used for the following step in the data processing workflow. c) Local resolution map for the 4.1 Å final map in b. d) Angular distribution for the particles leading to the 4.1 Å resolution map. e) Gold-standard Fourier shell correlation (FSC) curve for the final map.

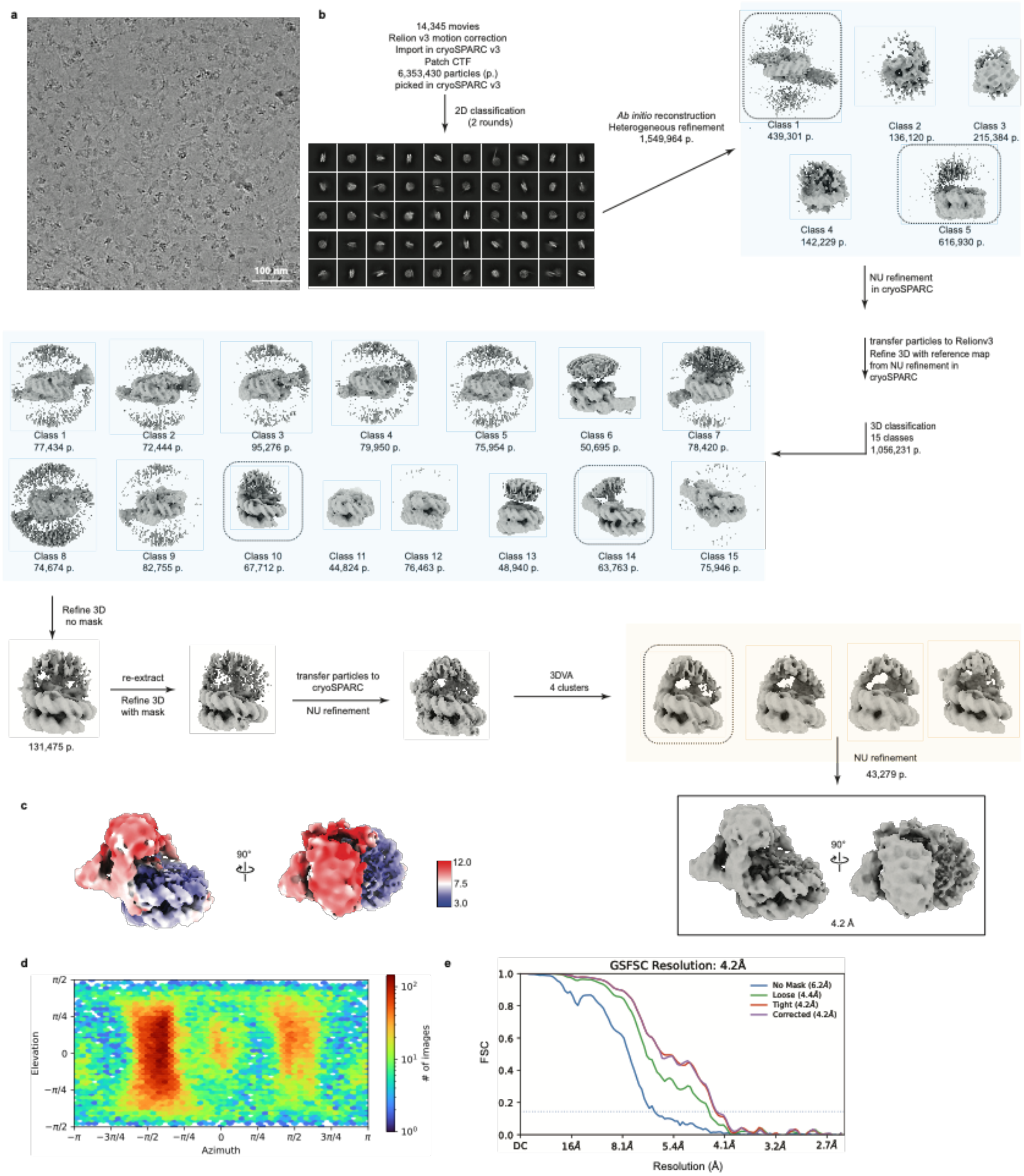

**Figure S3: cryo-EM data processing workflow for CLOCK:BMAL1-NCP$^{tandem\ 7bp}$.** a) Representative denoised cryo-EM micrograph. b) See methods. The movies were pre-processed within cryoFLARE. The resulting micrographs were imported in cryoSPARC v3 for particle picking. After multiple rounds of classification and refinements, the remaining particles were imported into Relion v3 for further classification and refinements. The final refinements and 3D variability analysis were performed in cryoSPARC v3. The final map is highlighted by a black box. The boxes defined by the dashed lines indicate the good maps and particle sets used for the following step in the data processing workflow. c) Local resolution map for the 4.2 Å final map in b. d) Angular distribution for the particles leading to the 4.2 Å resolution map. e) Gold-standard Fourier shell correlation (FSC) curve for the final map.

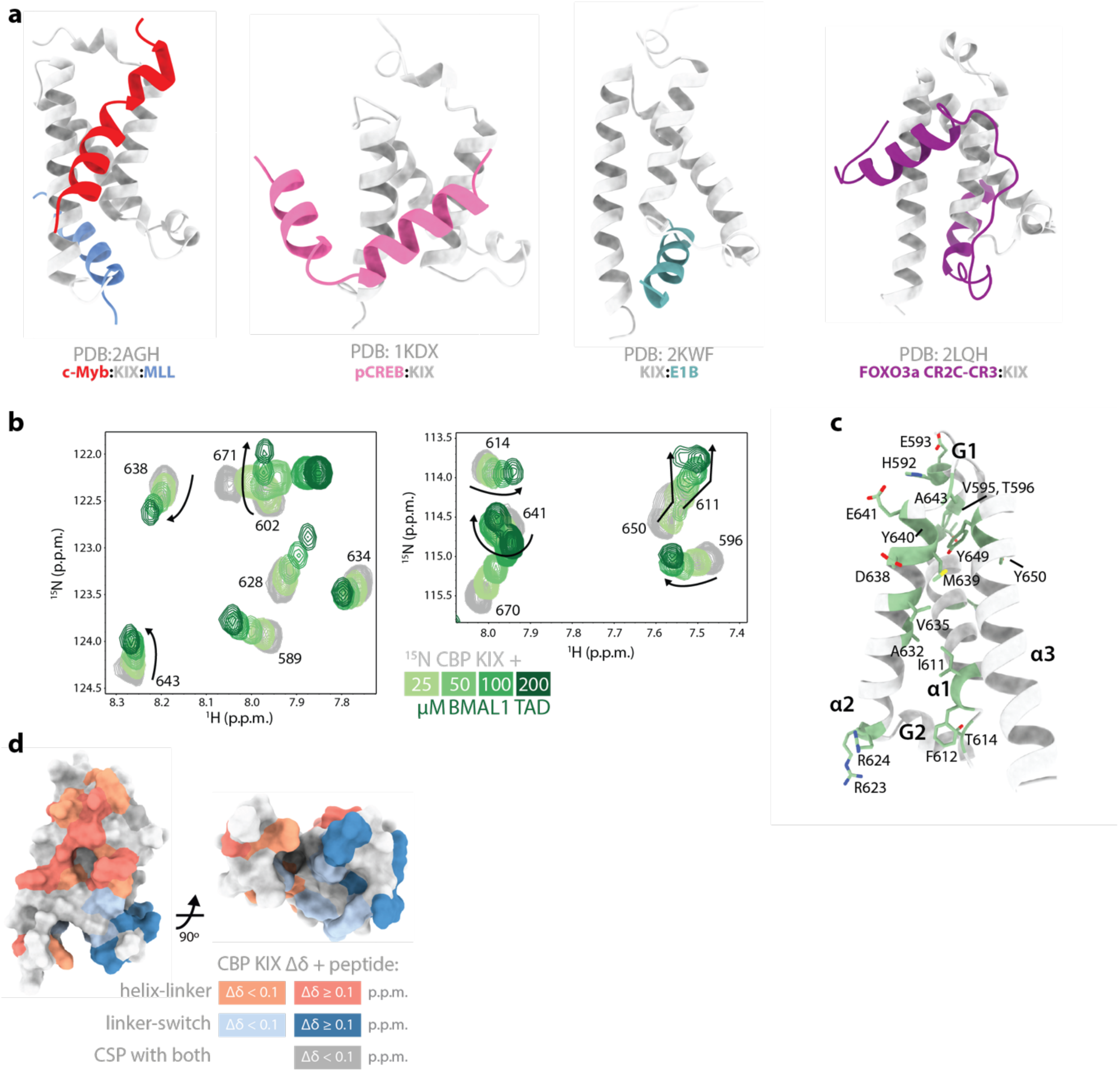

**Figure S4: Selected peaks in CBP KIX with curved CSP trajectories.** a) Structures of CBP KIX complexes with different transcription factors. b) Selected regions from the $^{15}$N-$^{1}$H HSQC of 100 μM $^{15}$N CBP KIX titrated with the BMAL1 TAD as indicated (from full spectrum shown in Fig. 2b). Chemical shifts with curved trajectories are marked with arrows. c) CBP KIX domain (PDB 2AGH, light gray) with secondary structure annotated and residues exhibiting curved chemical shift trajectories colored in green. For reference, the MLL1 binding site is located at the bottom of the KIX domain between the alpha-2 and alpha-3 helices beneath the G2 helix. d) CSPs of $^{15}$N CBP KIX bound to BMAL1 TAD peptides helix-linker (salmon hues) or linker-switch (steel blue hues) from Fig. 2e mapped onto PDB 2AGH. Residues that shift with both peptides are shown in dark gray. See Table 2 for more information.

**a**

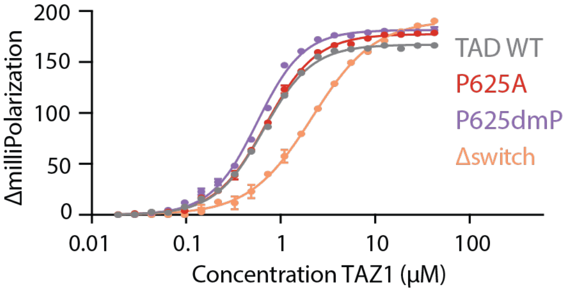

**b**

| | Kd (μM) | Hill coeff. |
|---|---|---|
| WT | 0.75 ± 0.14 | 1.59 ± 0.16 |
| P625A | 0.75 ± 0.05 | 1.55 ± 0.13 |
| P625dmP | 0.54 ± 0.12 | 1.71 ± 0.21 |
| Δswitch | 2.39 ± 0.23 | 1.12 ± 0.17 |

**Figure S5: CBP TAZ1 binding to variants of the BMAL1 TAD.** a) Fluorescence polarization data of CBP TAZ1 binding to different [5,6] TAMRA-BMAL1 minimal TAD probes as indicated: TAD WT (60% *trans,* 40% *cis*), P625A (all *trans)*, P625dmP (all *cis)*, Δswitch (truncated at F619Y) n = 3, mean ± s.d. b) $K_D$ and Hill coefficients from FP binding data.

**Table 1. Cryo-EM data collection, refinement and validation statistics**

| | Tandem 7bp<br>EMD-54869<br>PDB 9SFZ | Tandem 6bp<br>EMD-54868<br>PDB 9SFY |
|---|---|---|
| **Data collection and processing** | | |
| Detector | Falcon4i | Falcon 4i |
| Magnification voltage (kV) | 200 | 300 |
| Electron exposure ($e^{–}/Å^2$) | 50 | 50 |
| Defocus range (μm) | -0.8 - -2 | -0.8 - -2 |
| Pixel size (Å) | 0.84 | 0.845 |
| Symmetry imposed | C1 | C1 |
| Initial particle images (no.) | 6,353,430 | 6,840,426 |
| Final particle images (no.) | 43,279 | 24,238 |
| Map resolution (Å) | 4.2 | 4.1 |
| FSC threshold | 0.143 | 0.143 |
| Map resolution range (Å) | 3-12 | 3-12 |
| **Refinement** | | |
| Initial models used (PDB codes) | 6T93, 4F3L, 4H10, 8OSL | 6T93, 4F3L, 4H10, 8OSL |
| Model resolution (Å) | | |
| FSC threshold | 4.2 | 4.1 |
| Map sharpening *B* factor ($Å^2$) | N.A | N.A |
| Model composition | | |
| Non-hydrogen atoms | 13467 | 12309 |
| Protein residues | 960 | 861 |
| Nucleotides | 264 | 256 |
| Ligands | 0 | 0 |
| *B* factors ($Å^2$) | 193.80 | 144.61 |
| Protein | 179.69 | 128.88 |
| DNA | 214.81 | 165.77 |
| R.m.s. deviations | | |
| Bond lengths (Å) | 0.007 | 0.007 |
| Bond angles (°) | 0.867 | 0.931 |
| Validation | | |
| MolProbity score | 0.76 | 1.06 |
| Clashscore | 0.85 | 2.61 |
| Poor rotamers (%) | 0.16 | 0.95 |
| Ramachandran plot | | |
| Favored (%) | 99.04 | 97.98 |
| Allowed (%) | 0.96 | 1.90 |
| Disallowed (%) | 0.00 | 0.12 |
| **Model-to-data fit** | | |
| CCmask | 0.6571 | 0.6062 |
| CCbox | 0.7680 | 0.7276 |
| CCpeaks | 0.6071 | 0.5550 |
| CCvolume | 0.6474 | 0.5986 |

**Table 2. CBP KIX residues with CSPs mapped onto PDB 2AGH (chain 1.1/B).**

| Figure panel | Condition | CBP KIX residue |
|---|---|---|
| Fig. 2c | KIX with BMAL1 TAD, $\Delta\delta$ < 0.1 p.p.m. (light green) | 596,598,599,602,603,606,611,612, 620,621,622,624,625,628,631,632, 633,645,650,653,654,657,658,661, 662,664,668,670,671 |
| | KIX with BMAL1 TAD, $\Delta\delta$ > 0.1 p.p.m. (dark green) | 588,589,591,592,595,597,604,605, 610,614,616,626,627,629,634,635, 636,637,638,639,640,641,643,644, 646,649,652,655,672 |
| Fig. S4c | KIX with BMAL1 TAD, Curved CSP trajectories | 595,596,602,611,612,614,623,624, 632,635,638,639,640,641,643,649,671 |
| Fig. 2e Fig. S4d | KIX with TAD helix-linker $\Delta\delta$ < 0.1 p.p.m. (light salmon) | 590,594,603,605,642,645,647,656, 657,660,663,669 |
| | KIX with TAD helix-linker $\Delta\delta$ > 0.1 p.p.m. (salmon) | 602,606,608,646,648,650,654,658,670 |
| | KIX with TAD linker-switch $\Delta\delta$ < 0.1 p.p.m. (light blue) | 609,612,614,624,630,631,634,641,664 |
| | KIX with TAD linker-switch $\Delta\delta$ > 0.01 p.p.m. (blue) | 615,616,620,621,623,625,626,632, 635,636 |
| | KIX with helix-linker/linker-switch $\Delta\delta$ > 0.05 p.p.m. (dark gray) | 592,595,596,641 |

### Uncropped Blots

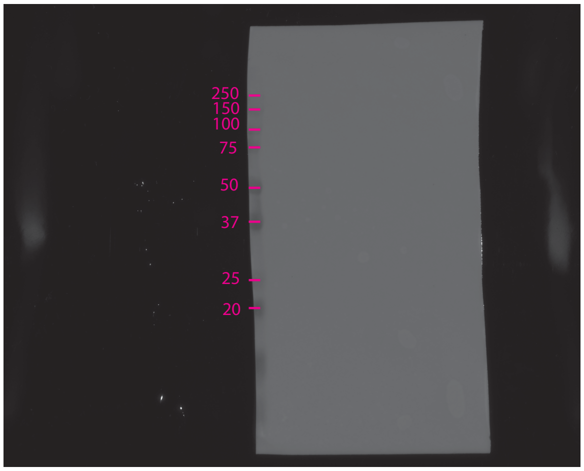

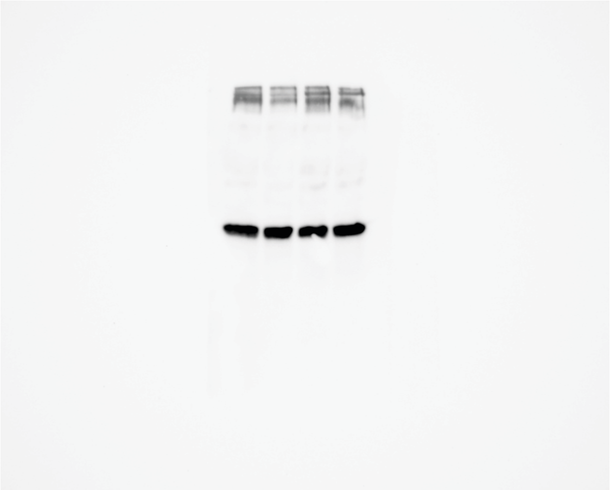
